# SCD1 and SCD5 Modulate PARP-Dependent DNA Repair via Fatty Acid Desaturation in Glioblastoma

**DOI:** 10.1101/2025.07.23.666454

**Authors:** Hayk Mnatsakanyan, Alessandro Sammarco, Rami Awwad, Abigail Hewett, Elie Roumieh, Caline Pechdimaljian, Yana Azar, Richa Pradhan, Baolong Su, Kevin J. Williams, Steven J. Bensinger, Christian E. Badr

## Abstract

Glioblastoma (GBM) relies on fatty acid metabolism to sustain its aggressive growth. While the role of stearoyl-CoA desaturase-1 (SCD1) in GBM is established, the function of its brain-enriched isoform, SCD5, remains unexplored. Here, we demonstrate that SCD5 is essential for glioblastoma stem cell (GSC) maintenance and genomic stability, with elevated expression in GSCs that declines upon differentiation, underscoring its role in tumor initiation. Through shotgun lipidomics, ^13^C metabolic flux analysis, and functional genomics, we reveal that SCD1 and SCD5 play non-redundant roles in fatty acid desaturation, with SCD5 preferentially desaturating C18:0 and uniquely contributing to sphingolipid remodeling. Genetic silencing of either isoform disrupts cell cycle progression, impairs DNA damage repair, and reduces GSC viability, while SCD5 knockdown significantly extends survival in orthotopic GBM models. Mechanistically, loss of SCD activity or saturated fatty acid accumulation triggers PARP1 hyperactivation and subsequent degradation, depleting RAD51 to compromise homologous recombination and induce parthanatos. These findings uncover a lipid-mediated vulnerability in GBM, linking fatty acid desaturation to PARP1-dependent genome integrity. Targeting SCD5 may offer a novel therapeutic strategy to eliminate therapy-resistant GSCs and enhance the efficacy of genotoxic or immunotherapeutic interventions.

## Introduction

Glioblastoma (GBM) remains the most aggressive and therapy-resistant primary brain tumor in adults. Despite standard treatment involving maximal surgical resection followed by radiation therapy (RT) and temozolomide (TMZ), which induce lethal DNA double-strand breaks, nearly all tumors recur due to intrinsic or acquired resistance^1^. This resistance is largely driven by active DNA damage repair (DDR) mechanisms^2,3^, which are particularly enhanced in glioblastoma stem-like cells (GSCs) ^4–6^. These observations underscore the need to target repair pathways in combination with conventional therapies.

Central to DDR is Poly (ADP-ribose) polymerase 1 (PARP1), which is overexpressed in various cancers including GBM ^7,8^. In GSCs, PARP1 plays a vital role in maintaining genomic stability and facilitating DNA repair, thereby enabling resistance to treatments such as RT and TMZ^9^. PARP1 also interacts with key DNA repair proteins such as RAD51^10,11^, which is essential for homology-directed repair (HDR), further contributing to GSC resilience^6^. Beyond its role in DDR, PARP1 is increasingly implicated in metabolic regulation, including lipid metabolism^12,13^, a critical driver of tumorigenesis, progression, and therapy resistance ^14,15^. Alterations in lipid metabolism and fatty acid biosynthesis, frequently observed in GBM ^16^, support rapid tumor growth and promote survival under therapeutic stress. Notably, upregulation of de novo lipogenesis is associated with poor clinical outcomes, and targeting lipid synthesis pathways has shown therapeutic promise in preclinical GBM models ^15–18^.

Fatty acid desaturation generates unsaturated fatty acids (UFAs), which serve diverse structural and signaling functions within cells ^19^. UFAs are incorporated into phospholipids ^20^, enhancing membrane fluidity, flexibility, and permeability, properties essential for maintaining cell integrity and supporting membrane trafficking and signal transduction ^20,21^. Stearoyl-CoA desaturase 1 (SCD1), the most abundant desaturase in humans, catalyzes the conversion of saturated long-chain fatty acids (SFAs) into monounsaturated fatty acids (MUFAs) ^22^, thereby promoting tumorigenesis and cancer progression in GBM ^17,23^. Our previous studies have shown that SCD1 is essential for GSC self-renewal and brain tumor initiation, with SCD1-targeted therapies demonstrating efficacy in GBM mouse models, particularly when combined with DNA-damaging agents like TMZ^17,23^.

A second desaturase, SCD5, remains largely understudied despite evidence pointing to alternative fatty acid desaturation pathways in GBM^18^ and in other cancers, such as liver and lung carcinoma ^24^. SCD5 is primarily expressed in the brain, ovary, and adrenal glands, though it is also present in other tissues ^25^. In this study, we establish that both SCD1 and SCD5 are critically required for GBM progression, functioning through overlapping and complementary mechanisms that regulate fatty acid desaturation, cell cycle progression, and DDR. These findings reveal a complex interplay between lipid metabolism and genomic maintenance and highlight new therapeutic avenues for targeting metabolic vulnerabilities in GBM.

## Results

### SCD5 is widely expressed across GBM subpopulations

Analysis of single-cell RNA sequencing (scRNA-seq) datasets from glioblastoma (GBM) patients^26,27^ revealed unexpectedly high expression of *SCD5* across multiple tumor subpopulations, whereas *SCD1* expression was confined to specific cellular clusters (Figure 1A–B and S1A). In contrast, bulk RNA-seq data from the GLASS consortium^28^ showed higher *SCD1* than *SCD5* expression in GBM (Figure 1C). This discrepancy likely reflects tumor heterogeneity, as bulk RNA-seq captures transcriptomic signals from both tumor and non-tumor components. A similar expression pattern was observed in normal brain tissue: scRNA-seq data showed predominant *SCD5* expression in neural cells, whereas bulk analyses favored *SCD1* (Figure 1D–E), suggesting that *SCD1* expression arises from a broader range of cell types beyond the neural lineage (Figure S1B).

**Figure 1.**
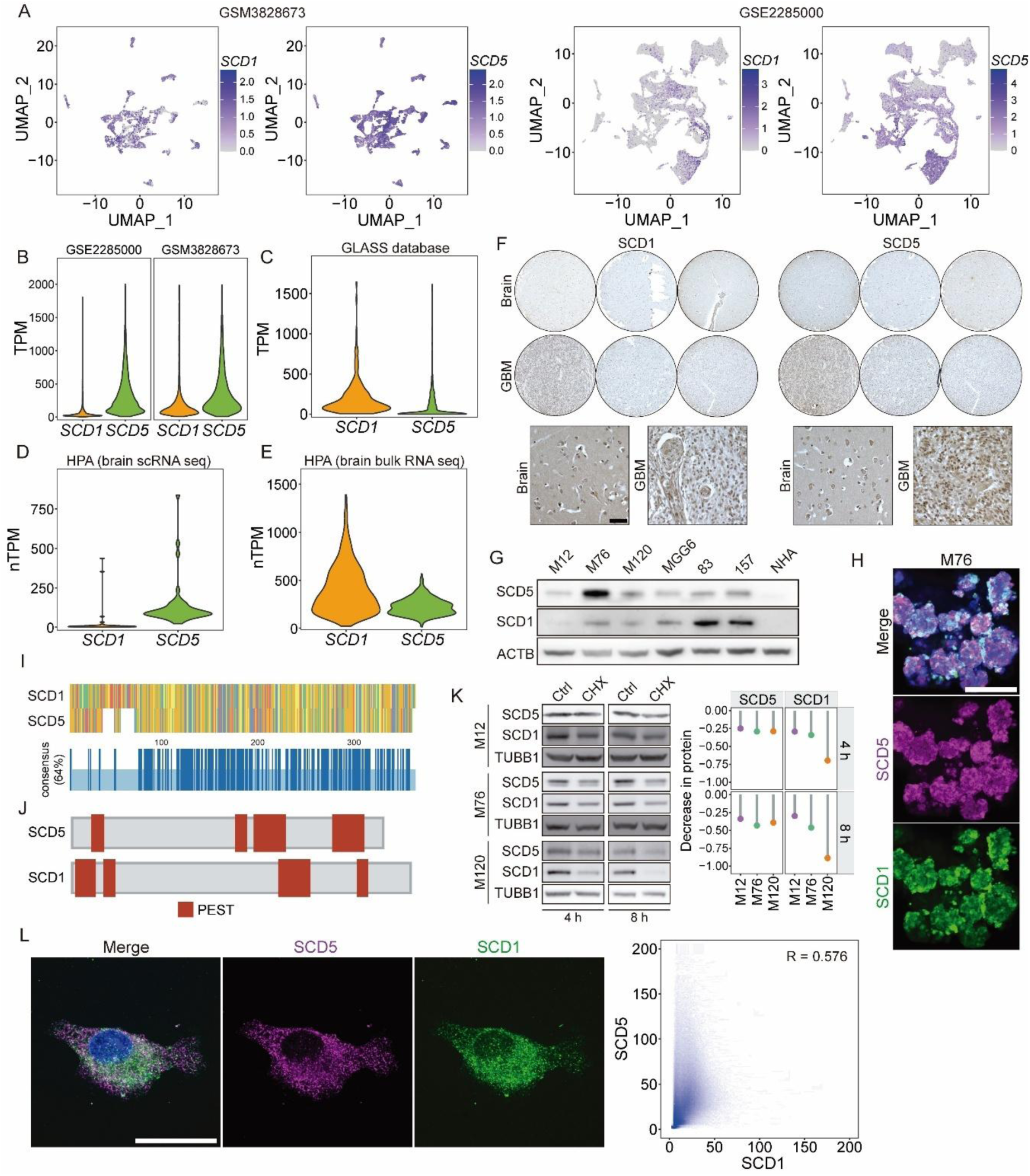
SCD5 is highly expressed in glioma cells at both transcriptional and translational levels. (A-B). scRNA seq analysis of two independent datasets (GSM3828673, n=28 patients; GSE2285000, n=39 patients) showing SCD1 and SCD5 expression patterns across cellular subpopulations. (A) (C) Bulk RNA-seq analysis of glioma tissues from GLASS consortium database comparing SCD1 and SCD5 expression. (D-E) scRNA seq (D) and bulk RNA-seq (E) analysis of normal brain tissue from Human Protein Atlas (HPA) database. (F) Immunohistochemical staining of SCD1 and SCD5 in healthy brain tissue and GBM patient tissue (n=3 each; top: overview; bottom: high-magnification insets). Scale bar = 50 µm. (G) Immunoblot analysis of SCD1 and SCD5 protein levels in six patient-derived GSC lines and NHA. (H) Immunostaining for SCD1 and SCD5 in GSCs. Scale bar = 50 µm. **I.** Protein sequence alignment of SCD1 and SCD5 with percentage amino acid similarity calculation. (J) Computational prediction of PEST motifs in SCD1 and SCD5 using ePESTfind. (K) Protein stability assay: Immunoblot and quantitative analysis of SCD1/SCD5 after cycloheximide treatment (100 µg/mL, 4-8 h) in three GSC lines. (L) Confocal microscopy analysis of SCD1/SCD5 co-localization with Pearson’s correlation coefficient (R) quantification. Scale bar = 25 µm.

To investigate SCD5 protein expression, we developed and validated a custom antibody. Specificity was confirmed through overexpression and knockdown experiments in GSCs using multiple shRNAs targeting both SCD5 transcript variants (Figure S1C-D), and further verified using miRFP670nano-tagged SCD1 and SCD5 constructs (Figure S1E-G). Immunohistochemical analysis demonstrated SCD5 protein expression in both GBM patient specimens and normal brain tissue (Figure 1F). Examination of seven patient-derived GSC lines representing diverse molecular subtypes revealed elevated protein expression of both SCD isoforms compared to normal human astrocytes (NHA) with distinct expression patterns between the two isoforms suggesting differential transcriptional or post-translational regulation (Figure 1G). Furthermore, co-immunostaining analysis demonstrated that SCD5 expression was uniform across GSCs, while SCD1 showed heterogeneous distribution, consistent with scRNA-seq findings (Figure 1H and S1J).

While SCD5 exists as two transcript variants (SCD5 and SCD5B), analysis of GLASS database samples and our GSC models revealed minimal expression of SCD5B (Figure S1H–I), suggesting this variant has limited biological relevance in GBM. Structural analysis showed that although SCD5 shares 65% amino acid similarity with SCD1 (Figure 1I), it exhibits distinct distributions of PEST motifs (Figure 1J), which are known to regulate the low stability of the SCD1 protein^29^. However, protein stability assays following cycloheximide (CHX) treatment demonstrated comparable degradation rates for both isoforms (Figure 1K). Subcellular localization studies confirmed that both enzymes reside in the endoplasmic reticulum (ER), but with minimal spatial overlap (Figures 1L and S1K–M), suggesting they occupy distinct functional microdomains. Together, these findings indicate that SCD5 is broadly expressed in GBM and, along with SCD1, likely contributes to tumor pathogenesis through complementary roles in lipid metabolism.

### Perturbation of SCD1 and SCD5 expression reveals distinct lipidomic profiles

To characterize the functional roles of SCD1 and SCD5 in lipid metabolism, we performed ^13^C metabolic flux analysis in two GSC lines following isoform-specific knockdown (KD). Both SCD1 and SCD5 KD resulted in reduced levels of newly synthesized MUFA (C16:1 and C18:1; Figure 2A). The levels of the PUFA C18:2 was undetected in the synthesized fraction, as expected (Figure 2A). However, we observed an increase in C18:2 levels in the total fraction, suggesting an increased PUFA uptake.

**Figure 2.**
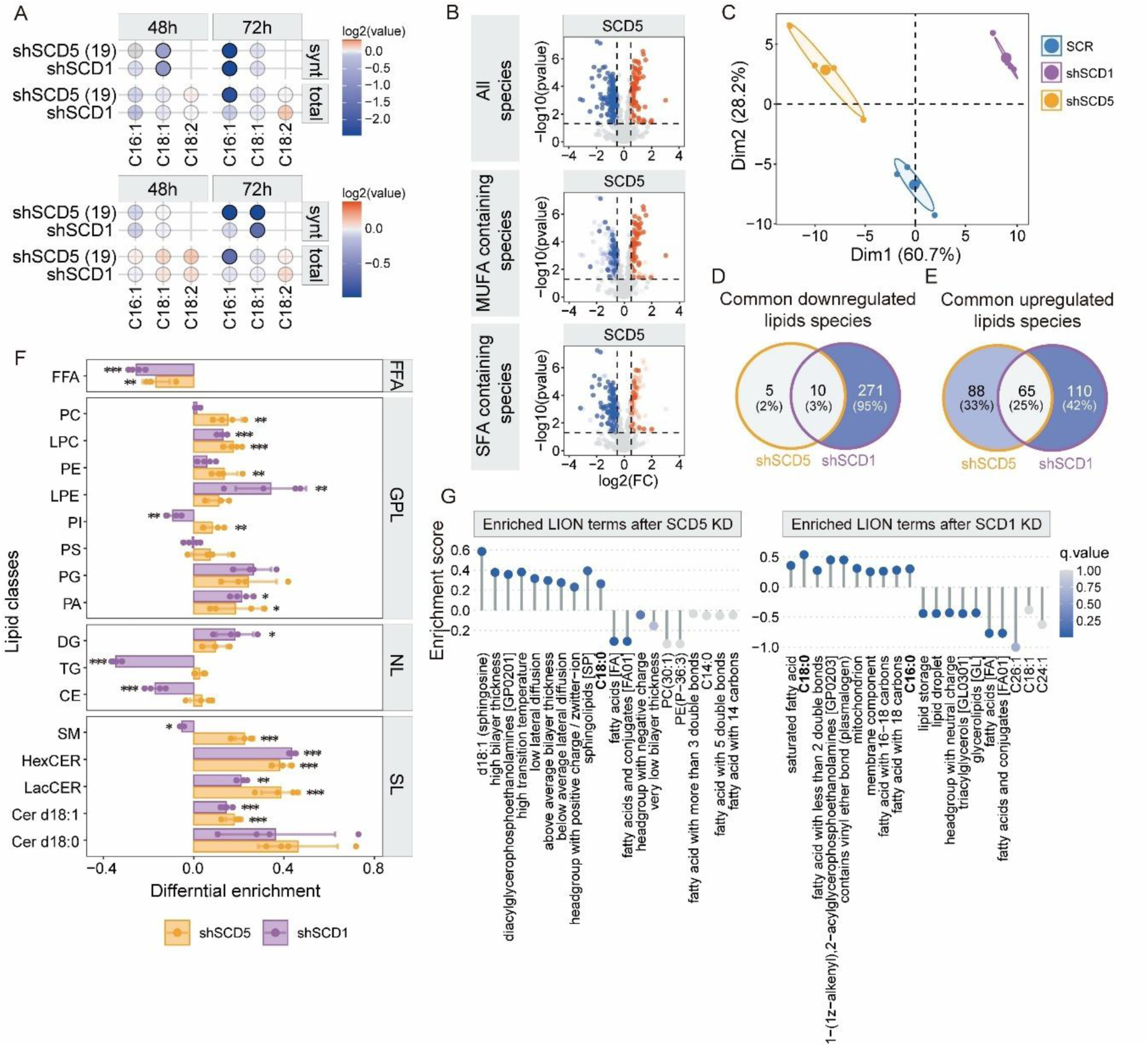
SCD1 and SCD5 silencing induce distinct lipidomic alterations in GSCs. (A) Isotope tracing analysis (¹³C incorporation) of FA synthesis pathways in GSCs following SCD1 or SCD5 knockdown (48-72 h). Dot plot displays log₂ fold changes in FA levels versus controls (filled dots: p < 0.05, Student’s t-test, n=4). (B) Volcano plot of lipid species alterations in SCD5-overexpressing GSCs versus controls (x-axis: log₂ fold change; y-axis: -log₁₀ p-value). (C) PCA of global lipid profiles following SCD1 or SCD5 knockdown. (D-E) Venn diagrams showing shared (D) downregulated and (E) upregulated lipid species between SCD1- and SCD5-deficient GSCs. (F) Differential abundance of lipid classes after 4-day SCD1/SCD5 silencing relative to SCR control (*p < 0.05, **p < 0.01, ***p < 0.001, Student’s t-test, n=4). (G) LION enrichment analysis of top 10 significantly altered biological terms.

To comprehensively characterize the lipidomic changes mediated by SCD1 and SCD5 and elucidate their non-redundant functions, we employed shotgun lipidomics as an unbiased discovery approach. First, we confirmed the desaturase activity of SCD5 in GSCs through overexpression studies, which demonstrated a significant increase in MUFA and a corresponding decrease in SFA (Figure 2B). Subsequent lipidomic profiling of SCD1- and SCD5-KD GSCs revealed distinct lipid remodeling patterns. Principal component analysis (PCA) separated the lipidomic profiles of SCD1-KD and SCD5-KD cells from controls and from each other, indicating isoform-specific metabolic roles (Figure 2C and S2A). While SCD1 knockdown induced broad alterations across multiple lipid classes, SCD5 knockdown resulted in more selective changes, with only 15 lipid species showing significant downregulation (Figures 2D and S2B). Intriguingly, both knockdown conditions led to increased levels of several MUFA-containing lipids, suggesting compensatory activation of the remaining SCD isoform (Figure S2B). A potential exogenous source of these lipids can be ruled out, as the cell culture medium lacked C18:1.

Comparative analysis of affected lipid species revealed that ∼74% of upregulated lipids following SCD5 KD were distinct when compared to SCD1 KD (Figures 2E and S2C), supporting non-redundant functions of these enzymes. Lipid classes analysis showed that both KDs decreased free fatty acids, while differentially affecting other lipid classes (Figure 2F). SCD5 KD increased most glycerophospholipids (Figure S2D) and upregulated sphingolipids, particularly sphingomyelin with saturated acyl chains (Figure 2F and S2E). In contrast, SCD1 KD increased diacylglycerols while decreasing cholesterol esters and triacylglycerides, leading to reduced lipid droplet formation (Figures 2F and S2F).

To functionally interpret the lipidomic alterations, we performed Lipid Ontology (LION) enrichment analysis ^30^, which links changes in lipid species to their biophysical and functional characteristics. In SCD5-KD GSCs, the most significantly upregulated terms were related to membrane characteristics, including increased bilayer thickness, elevated transition temperature, and reduced lateral diffusion. The most downregulated terms were associated with free fatty acid (FFA) abundance (Figure 2G). In contrast, SCD1 KD primarily enriched terms related to lipid saturation and significantly reduced those linked to lipid storage, lipid droplets, and neutral lipid content, consistent with the observed decrease in lipid droplet formation (Figures S2F). At the molecular level, SCD5 KD was associated with enrichment of ceramide- and phosphatidylethanolamine-related terms, whereas SCD1 KD was linked to an increase in plasmalogens (acylglycerophosphoethanolamines and lipids with vinyl ether bonds) and mitochondrial membrane components. Importantly, several of these lipid remodeling patterns were reversed in SCD5-overexpressing cells (Figure S2G). Notably, LION analysis revealed an increase in SFAs, particularly C18:0, following both SCD1 and SCD5 knockdown (Figure 2G). However, while SCD1 knockdown also led to an increase in C16:0, this was not observed in SCD5-deficient cells. Supporting these findings, SCD5 overexpression resulted in a decrease in C18:0 and a corresponding increase in C18:1 levels (Figure S2G).

Overall, our findings demonstrate two key functional distinctions: 1) SCD5 appears specialized for C18:0 to C18:1 conversion, while SCD1 desaturates both C16:0 and C18:0; and 2) SCD5 KD uniquely increases sphingolipids, particularly saturated sphingomyelins, without affecting neutral lipids, whereas SCD1 KD alters neutral lipid metabolism and lipid droplet formation. These differences, coupled with their distinct ER localizations, suggest that SCD1 and SCD5 operate in separate metabolic networks despite sharing desaturase activity.

### SCD5 preferentially desaturates C18:0 but not C16:0

Our results suggest that SCD1 and SCD5 exhibit differential substrate specificity. To test this, we analyzed FA composition after SCD1 or SCD5 KD (Figure S2A) or SCD5 overexpression (Figure S2B). Shotgun lipidomics revealed distinct FA profiles between these conditions. Notably, while SCD1 KD significantly reduced the desaturation index (DI) of C18:0, SCD5 KD showed no significant effect on the DI of either C16:0 or C18:0 (Figure 3A), suggesting potential functional compensation by SCD1. Importantly, SCD5 overexpression specifically enhanced the DI of C18:0 without affecting C16:0 DI (Figure 3B), providing further evidence that SCD5 selectively desaturates C18:0.

**Figure 3.**
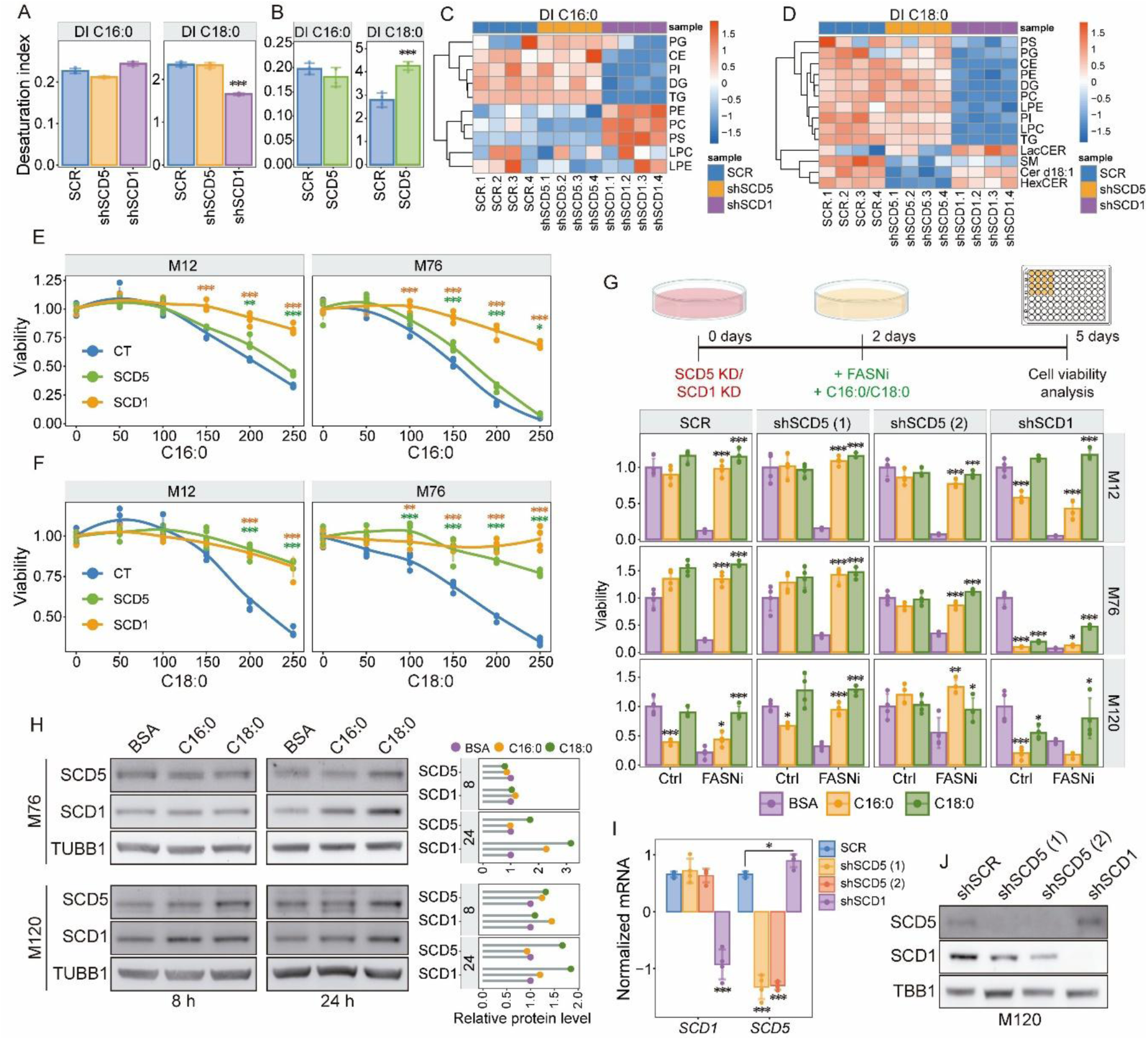
SCD5 preferentially desaturates C18:0 in GSCs. (A) Desaturation indices (C16:1/C16:0 and C18:1/C18:0) in GSCs following 4-day SCD1 or SCD5 knockdown (mean ± SD; *p<0.05, **p<0.001, ***p<0.0001, Student’s t-test, n=4). (B) Desaturation indices in SCD5-overexpressing GSCs. (C-D) Heatmaps of lipid class-specific desaturation indices for (C) C16:0 and (D) C18:0 following SCD1/SCD5 knockdown. (E-F) Dose-response viability curves of GSCs overexpressing SCD1 or SCD5 treated with (E) C16:0 or (F) C18:0 for 4 days (mean ± SD, n=4). (G) Three GSCs lines were treated with FASNi ± C16:0/C18:0, 2 days after SCD1/SCD5-knockdown. Cell viability was measured 3 days after treatment and represented as fold change relative to BSA/DMSO-treated control (bar plot: mean ± SD; *p<0.05, **p<0.001, ***p<0.0001, n=4). (H) Immunoblot analysis of SCD1/SCD5 in GSCs treated with C16:0 or C18:0 for 8 or 24 h. (I) qPCR analysis of SCD1/SCD5 mRNA 3 days post-knockdown in four GSC lines (M12, M76, 83, and M120). ((J) Immunoblot analysis of SCD1/SCD5 protein levels 3 days following knockdown.

To dissect these roles further, we calculated DIs across lipid classes (Figures 3C–D, S3C). SCD1 KD reduced the DI of C16:0 in some lipid species but increased it in others, resulting in no net change in the global lipidome. In contrast, SCD1 KD consistently decreased the DI of C18:0 across most lipid classes, except sphingolipids (SLs; e.g., LacCER, Cer d18:1, HexCER). Strikingly, SCD5 KD only reduced the DI of C18:0 in SLs (Cer d18:1, HexCER, SM), underscoring its role in SL metabolism.

Functional assays corroborated substrate specificity. Overexpression of SCD1 protected GSCs from lipotoxicity induced by both C16:0 and C18:0, whereas SCD5 overexpression only rescued C18:0 toxicity in two GSC lines (Figures 3E–F). To corroborate these findings, we silenced either SCD1 or SCD5, inhibited de novo FA synthesis using fatty acid synthase inhibitors (FASNi), and attempted to rescue GSC viability by supplementing the culture medium with either C16:0 or C18:0 (Figure 3G). In GSCs expressing a non-targeting shRNA (SCR), both C16:0 and C18:0 rescued cell viability after FASN inhibition (Figure 3G). In SCD5 KD cells, adding either C16:0 or C18:0 prevented the loss of cell viability due to FASNi, since SCD1 expression remained unaffected and could desaturate both C16:0 and C18:0 to produce functional lipids. However, after SCD1 KD, adding C16:0 led to a significant increase in cell toxicity, even in GSCs not exposed to FASNi (Figure 3G). Conversely, C18:0 supplementation in SCD1 KD cells treated with FASNi rescued cell viability (Figure 3G), further supporting an exclusive role of SCD5 in C18:0 desaturation.

SFA supplementation increases SCD1 expression^23,31^. Further, we have previously shown that the pharmacological inhibition of SCD results in increased SCD1 protein levels, due to SFA accumulation^23^ . Consistently, treatment with both C16:0 and C18:0 upregulated SCD1 expression in both GSC lines (Figure 3H). In contrast, only C18:0 induced SCD5 expression in both lines (Figure 3H). Additionally, SCD1 knockdown in four GSC lines triggered a transient compensatory upregulation of SCD5, which was only detectable at early time points (Figures 3I-J). This asymmetry is likely because SCD1 knockdown significantly reduces the DI of C18:0 across most of the lipidome, while SCD5 knockdown does not cause widespread changes beyond the SL class.

In sum, these data demonstrate that SCD5 is a dedicated C18:0 desaturase with a pronounced role in SL metabolism, while SCD1 exhibits broader activity toward both C16:0 and C18:0.

### SCD5 is essential for GSC self-renewal, survival, and tumor growth

To explore whether SCD expression is linked to GSC stemness, we analyzed microarray data comparing different GBM cell types (GDS3885)^32^, including differentiated GBM cell lines (cultured in serum), GSCs grown as neurospheres, and primary GBM cells (Figure 4A). Correlation analysis revealed a positive correlation between *SCD5* and markers of neural stem and progenitor cells (Figure 4B). We observed that cells cultured in serum (differentiation conditions) lost the expression of *SCD5* along with stemness markers like *SOX2*, *NES*, *OLIG1*, *PROM1* (CD133), and *OLIG2*. In contrast, non-primary cell lines, regardless of whether they are cultured with serum or serum-free conditions, exhibited increased *SCD1* expression. These findings highlight key metabolic shifts, including the loss of *SCD5* and upregulation of *SCD1* in established cell lines, which do not recapitulate the SCD expression patterns observed in patient tumors or primary GSCs.

**Figure 4.**
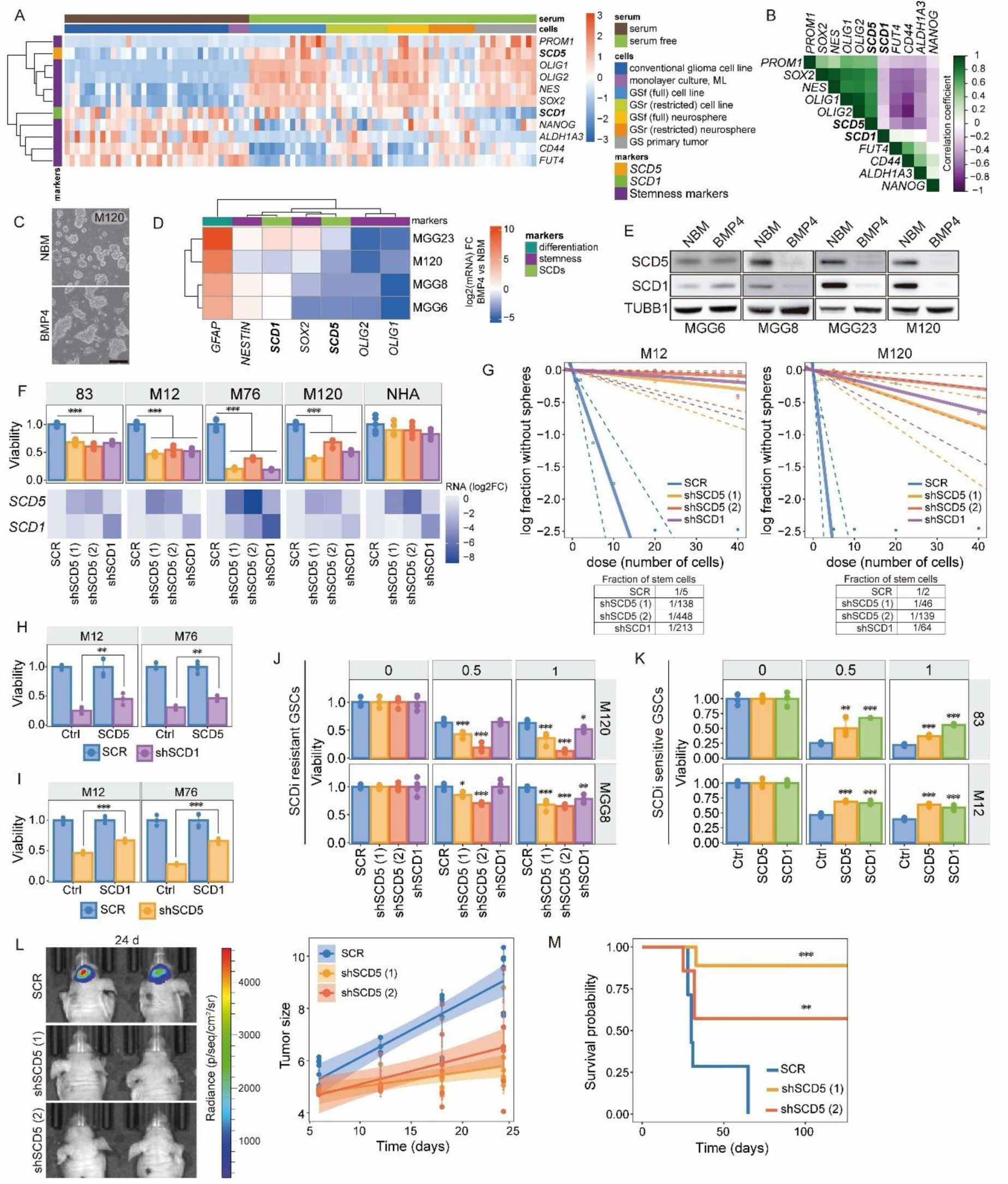
SCD1 and SCD5 are essential for GSC maintenance and tumorigenesis. (A) Heatmap of SCD5, SCD1, and stemness marker expression across glioma cell lines, neurosphere-cultured GSCs, and primary GSCs (GDS3885 microarray dataset). (B) Spearman correlation analysis between SCD isoforms and stemness markers using the same transcriptomic data. (C) Morphological changes in GSCs following 6-day BMP4 differentiation (20 ng/mL). Scale bar: 100 µm. (D) Heatmap of stemness (NES, OLIG1/2, SOX2) and differentiation (GFAP) marker expression post-BMP4 treatment. (E) Immunoblot analysis of SCD1/SCD5 protein levels after BMP4-induced differentiation. (F) Viability assay of GSCs following 4-day SCD1/SCD5 knockdown (mean ± SD; n=4; *p<0.05, **p<0.001, ***p<0.0001, Student’s t-test). Lower panel: mRNA knockdown efficiency heatmap. (G) ELDA of stem cell frequency in M12/M120 GSCs after SCD1/SCD5 knockdown (14 days). (H-I) Rescue experiments: Viability of (H) SCD1-overexpressing GSCs with SCD5 knockdown and (I) SCD5-overexpressing GSCs with SCD1 knockdown (n=4). (J-K) Drug response assays: (J) SCD1i-resistant GSCs (treated with the SCD1 inhibitor CAY10566, 1 µM) with SCD5 knockdown, and (K) SCD1i-sensitive GSCs overexpressing SCD5 (n = 4). (L) In vivo tumor growth measured by Fluc bioluminescence imaging of control (SCR) vs. SCD5-knockdown GSCs (n=7 mice/group) at indicated time points. (M) Kaplan-Meier survival curves of mice bearing control or SCD5-deficient GSCs (n=7/group).

Further supporting this, analysis of the DepMap database showed that established GBM cell lines (grown in serum) depend on SCD1 but not SCD5 (Figure S4A). To validate these findings, we differentiated patient-derived GSCs using BMP4 (Figures 4C-E) and assessed the mRNA and protein levels of stemness and differentiation markers, along with SCD1 and SCD5. Upon differentiation, *SCD5* mRNA levels decreased in all GSC lines tested (Figure 4D), while protein levels of both SCD1 and SCD5 were markedly reduced in three of four lines (Figure 4E). These results align with the microarray data, confirming that SCD5 is associated with stemness in GSCs. In contrast, while SCD1 levels declined during BMP4-induced differentiation, its upregulation in long-term serum culture suggests distinct regulatory mechanisms. This likely reflects the necessary role of SCD1 in maintaining basal lipid homeostasis and membrane integrity in rapidly dividing cells while also supporting GSC self-renewal through mechanisms independent of stemness regulation.

To assess whether SCD5, like SCD1, is essential for GSC self-renewal, we performed shRNA-mediated KD in multiple GSC lines and NHA cultures. KD of either SCD1 or SCD5 minimally affected NHAs but significantly reduced GSC viability (Figure 4F). Extreme limiting dilution assays (ELDA) and secondary sphere formation assays further demonstrated impaired stem cell frequency and sphere-forming capacity (Figures 4G and S4B-C), confirming that both enzymes are required for GSC self-renewal. To rule out off-target effects, we engineered a lentiviral vector expressing either a wild-type SCD5 (SCD5wt) or an shSCD5-resistant SCD5 (SCD5^shRes^; resistant to shSCD5 (2)) fused to miRFP670nano to monitor protein stability post-KD (Figures S4D-E). GSCs expressing miRFP670nano-SCD5^shRes^ maintained fluorescence and viability upon shSCD5 (2) transduction, whereas shSCD5 (1) reduced both (Figures S4F-G).

To determine if the viability loss was due to the metabolic function of SCD5, we supplemented GSC medium with C18:1 (the direct product of SCD), post-KD. This significantly rescued the viability loss in all four GSC lines after SCD1 or SCD5 KD (Figure S4H). To test functional redundancy, we overexpressed SCD5 in SCD1-KD cells (and vice versa) and found only partial rescue (Figure 4H-I), supporting our lipidomic data that SCD1 and SCD5 have distinct biological roles despite shared enzymatic activity.

Resistance to SCD inhibitors (SCDi) has been associated with alternative desaturation pathways; however, the role of SCD5 in this context had not been previously explored ^18,24^. Using a panel of SCDi-sensitive and -resistant GSCs we previously characterized^17^, we found that SCD5 knockdown sensitized cells to pharmacological SCD1 inhibition with CAY10566 (Figure 4J), whereas overexpression of either SCD1 or SCD5 conferred resistance (Figure 4K). These findings suggest that SCD5 functions as an alternative source of MUFAs, compensating for SCD1 inhibition.

Finally, to assess the role of SCD5 in tumorigenesis, we intracranially implanted GSCs expressing either SCR or two shRNAs targeting SCD5 into the brains of nude mice. SCD5 KD significantly suppressed tumor growth and extended survival (Figures 4L–M) with 7/8 mice in the shSCD5 (1) group and 4/7 in the shSCD5 (2) group surviving to 240 days, with no detectable tumors by bioluminescence imaging (Figure S4I). Together, these findings establish SCD5 as a critical regulator of GSC maintenance and a potential therapeutic target in GBM.

### SCD activity is necessary for cell cycle progression

To identify gene expression changes and regulatory networks affected by SCD loss, we performed bulk RNA-seq on GSCs following SCD1 or SCD5 KD (Figure 5A). Gene set enrichment analysis (GSEA) revealed that the top five significantly downregulated biological processes were shared between SCD1 and SCD5 depletion, underscoring functional redundancy between these two enzymes (Figure 5B).

**Figure 5.**
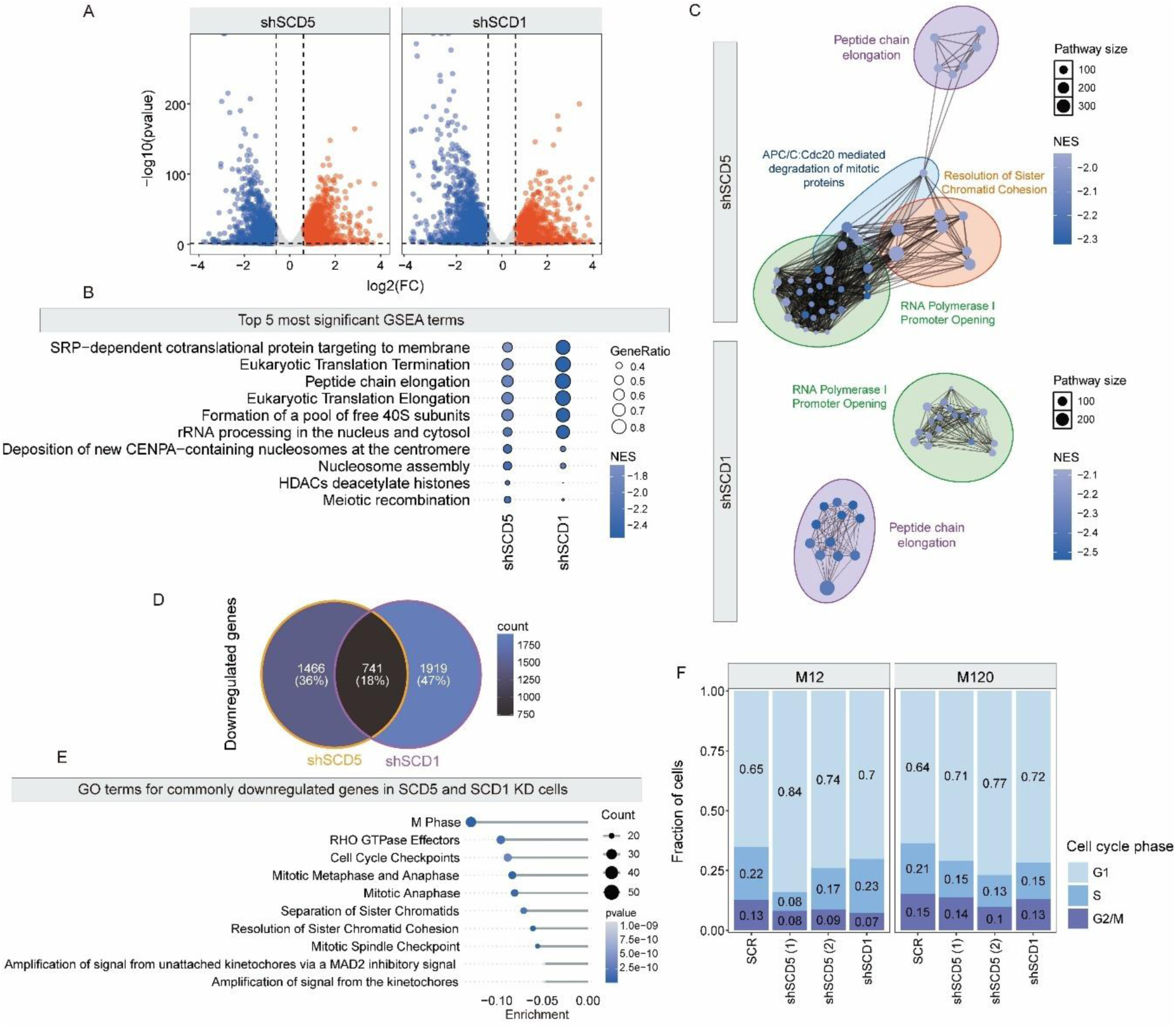
SCD1 and SCD5 regulate cell cycle progression in GSCs. (A) Transcriptomic changes in GSCs following 4-day SCD1 or SCD5 knockdown (volcano plot: x-axis = log₂ fold change; y-axis = -log₁₀ p-value). (B) Top 5 significantly enriched GO terms from gseGO analysis. Terms are ranked by lowest adjusted *p*-values (dot size = gene count; color = adjusted p-value; filled dots: p.adj<0.05, n=4). (C) Enrichment map of 40 most altered GO terms. (D) Venn diagram of downregulated transcripts shared between SCD1- and SCD5-deficient GSCs. (E) GO enrichment analysis of commonly downregulated genes in SCD1/SCD5 knockdown conditions. (F) Cell cycle distribution analysis by flow cytometry 4 days post-SCD1/SCD5 knockdown.

Visualization of the top 30 significantly altered pathways in an enrichment map further demonstrated that clustered terms were predominantly associated with cell cycle progression, mRNA translation, and protein synthesis (Figure 5C). Consistent with this, Gene Ontology (GO) analysis showed that the top 10 most downregulated terms after SCD1/SCD5 KD were linked to cell division (Figure 5E). Conversely, upregulated terms included neurotrophic signaling and responses to metal ions (Figure S5A-B), with the former potentially reflecting a compensatory mechanism by which GSCs foster adaptative neuron-cancer cell interactions^33^ to survive SCD depletion.

To functionally validate these observations, we analyzed cell cycle distribution using flow cytometry four days post-KD. Depletion of either SCD1 or SCD5 resulted in G1-phase accumulation and a concomitant reduction in G2/M phase cells (Figure 5F), mirroring the transcriptional changes. Furthermore, synchronized GSCs (double-thymidine block) released into the cell cycle exhibited delayed progression at 3, 6, and 9 hours post-release upon SCD KD, compared to controls (Figure S5C). Collectively, these data demonstrate that SCD activity is essential for efficient cell cycle progression in GSCs.

### SCD downregulation triggers PARP hyperactivation and parthanatos

While the mechanistic details remain unclear, our previous work demonstrated that SCD inhibition depletes RAD51, impairs HDR, and increases DNA damage in GSCs ^17,23^. RNA-seq analysis revealed significant downregulation of base excision repair pathways following SCD1 or SCD5 KD (Figure 6A), corroborated by increased DNA damage evident through elevated γ-H2A.X levels (Figure 6B and S6A).

**Figure 6.**
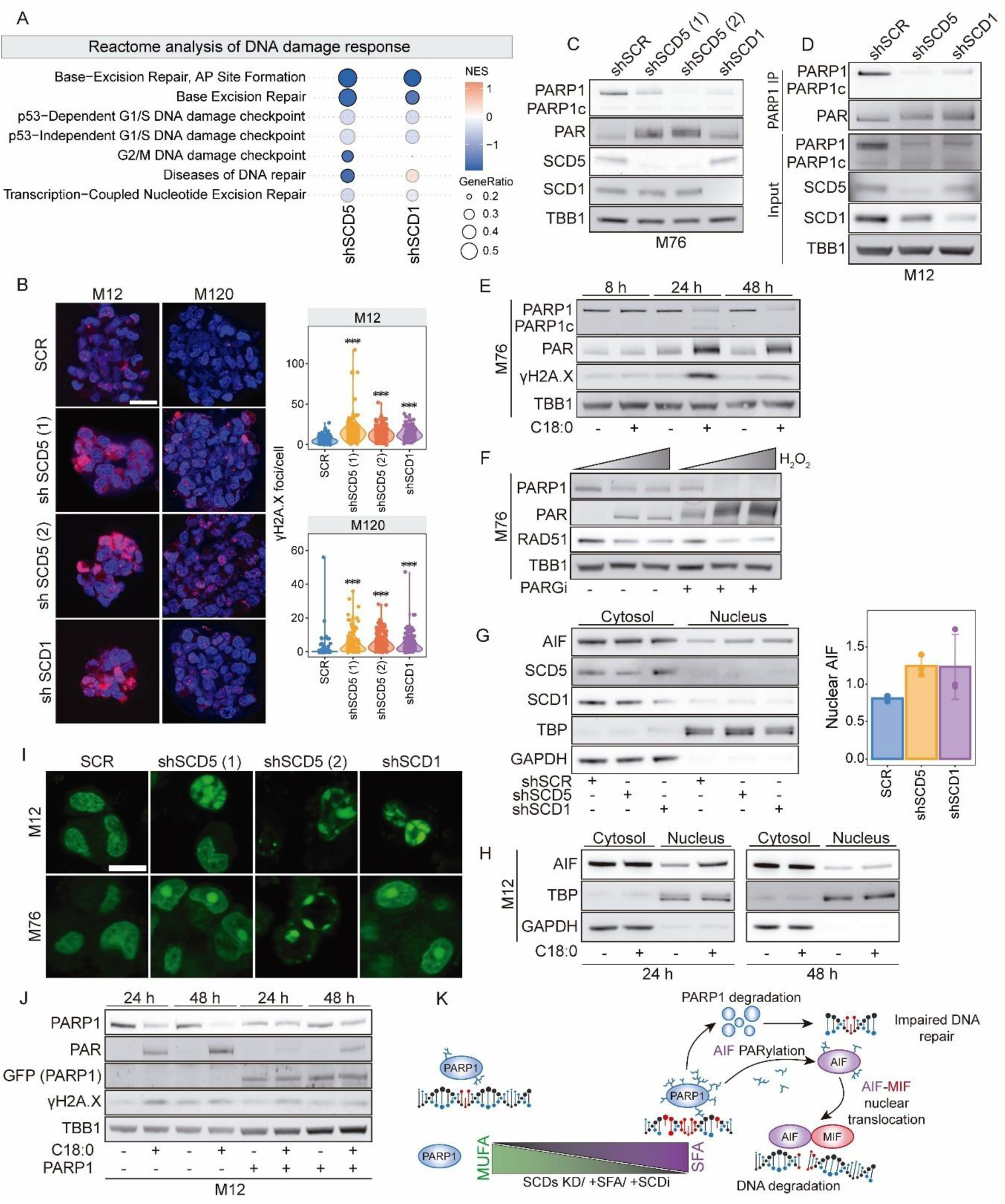
SFA accumulation induces DNA damage, PARP1 hyperactivation, and parthanatos in GSCs. (A) Reactome pathway analysis of DNA damage repair mechanisms altered by SCD1/SCD5 knockdown (4 days). (B) γH2AX immunofluorescence (left) and quantification (violin plot, right) in SCD1/SCD5-deficient GSCs (n>100 cells; *p<0.05, **p<0.001, ***p<0.0001, Student’s t-test). Scale bar: 25 µm. (C) Immunoblot analysis of PARP1 and PARylation at the PARP1 expected molecular weight post-SCD1/SCD5 knockdown. (D) PARP1 immunoprecipitation with PAR immunoblot in knockdown conditions. (E) Time-course immunoblot analysis of PARP1, PAR, and γH2AX in GSCs treated with 150 µM of C18:0. (F) immunoblot analysis of PARP1, PAR, and RAD51 expression after treatment with H₂O₂ (0-2 mM, 15 min) and PARG inhibitor (1 µM, 24 hr). (G-H) Subcellular fractionation followed by immunoblot analysis of AIF expression following SCD1/SCD5 knockdown (G) or C18:0 treatment (H). TATA-binding protein (TBP; nuclear) and GAPDH (cytoplasmic) were used as loading controls. Quantification shows mean nuclear AIF/TBP ratio ± SD (n=3 GSC lines: M12, M76, M120). (I) PicoGreen staining of nuclear DNA fragmentation post-SCD1/SCD5 knockdown. Scale bar: 10 µm. (J) Immunoblot analysis of DNA damage markers in PARP1-GFP–expressing GSCs treated with 150 µM of C18:0 for 24 and 48h. (K) Schematic illustration of the proposed mechanism by which saturated fatty acid (SFA) accumulation induces DNA damage, leading to PARP1 hyperPARylation, subsequent PARP1 degradation, and parthanatos.

PARP1, the predominant PARP isoform in GBM (Figures S6B), plays a central role in DNA repair, including BER and HDR ^34^. Prior studies have established that PARP inhibition transcriptionally represses RAD51^35^, implicating PARP1 in the regulation of DNA repair machinery. Although PARP1 mRNA levels were unchanged post-SCD KD (Figure S6C), we observed reduced PARP1 protein without caspase-mediated cleavage (no increase in PARP1c). Strikingly, autoPARylation, detected by PAR bands at PARP1 molecular weight, was markedly enhanced (Figure 6C), confirmed by PARP1 immunoprecipitation (Figure 6D). Conversely, SCD1 or SCD5 overexpression reduced γ-H2A.X and restored PARP1 levels (Figure S6D), while BMP4-induced differentiation (which lowers SCD expression) mirrored SCD KD effects (Figure S6E). Further, we observed a robust inverse correlation between PARP1 and γ-H2A.X across GSC lines (Figures S6F-G), suggesting a mechanistic link between SCD, PARP1 and γ-H2A.X.

To determine whether saturated fatty acid (SFA) accumulation mediates the observed effects, we supplemented GSCs with C18:0, a direct substrate of SCD. This treatment triggered a time-dependent cascade characterized by PARP1 hyper-PARylation, subsequent depletion of PARP1 protein, and increased DNA damage, all of which peaked at 24 hours before partially recovering—likely due to the activation of compensatory DNA repair mechanisms (Figures 6E, S6H). Notably, RAD51 levels declined markedly at 48 hours, coinciding with the lowest observed PARP1 levels (Figure S6H). Further analysis revealed that the C18:0-induced DNA damage was mediated by reactive oxygen species (ROS), as co-treatment with the antioxidant N-acetylcysteine (NAC) significantly reduced γ-H2A.X expression (Figure S6I). These findings suggest that sustained PARP1 auto-PARylation leads to its degradation, which in turn contributes to RAD51 depletion.

To directly test this hypothesis, we treated GSCs with hydrogen peroxide (H₂O₂) to induce PARP1 auto-PARylation, in combination with a PARG inhibitor (PARGi) to prevent PAR removal. This dual treatment led to a synergistic reduction in PARP1 protein levels (Figures 6F and S6J-K), confirming that persistent PARylation promotes PARP1 degradation.

We next investigated the degradation mechanisms by treating GSCs with C18:0 for 48 hours, followed by administration of either autophagy inhibitors (Bafilomycin A or 3-Methyladenine) or proteasome inhibitors (MG-132 or Bortezomib). Both treatments restored PARP1 protein levels (Figure S6L), indicating that PARP1 is degraded via both autophagy and proteasomal pathways. Consistent with this, we observed increased LC3 cleavage following C18:0 treatment, SCD inhibition, or SCD knockdown (Figure S6M), demonstrating that SFA accumulation activates autophagy and contributes to PARP1 degradation.

Excessive PARP1 autoPARylation can induce parthanatos, a form of cell death where nuclear PAR translocates to the cytosol, binding apoptosis-inducing factor (AIF) and triggering its nuclear import ^36^. SCD KD or C18:0 treatment increased nuclear AIF (Figures 6G-H, S6N-P) and cytosolic PAR accumulation (Figure S6N). Once in the nucleus, AIF recruits migration inhibitory factor (MIF), which cleaves DNA into large fragments, resulting in chromatin condensation and fragmentation^36^. PicoGreen staining revealed chromatin condensation and fragmentation in SCD-depleted GSCs (Figure 6I), hallmarks of parthanatos. Elevated PARP1 levels allow cells to repair DNA without excessive accumulation of PARylated PARP, thereby preventing parthanatos^37^. Crucially, PARP1 overexpression mitigated DNA damage, autoPARylation, and cell death (Figures 6J, S6Q-R), confirming that PARP1 levels determine cell fate under SFA stress.

In conclusion, SCD deficiency or SFA accumulation hyperactivates PARP1, leading to its autodegradation via autophagy/proteasomal pathways, impaired DNA repair, and parthanatos (Figure 6K).

## Discussion

Our study establishes SCD5 as a critical metabolic regulator in GBM, with dual functions in maintaining GSC self-renewal and modulating DNA damage repair pathways. We demonstrate that SCD5 is essential for GSC tumor-initiating capacity, and we reveal previously unrecognized mechanistic links between fatty acid desaturation and genomic stability in GBM.

While the role of SCD1 in cancer is well established, the function of SCD5 remains poorly understood. Conflicting reports have described SCD5 depletion as having no effect on proliferation ^18,38^, promoting necrosis ^39^, or even enhancing cell growth ^40^. Similarly, ectopic expression of SCD5 has been linked to both pro-tumorigenic effects, such as promoting thymic cancer cell migration ^41^, and anti-tumorigenic outcomes, including reduced metastasis in breast cancer and melanoma models^42,43^. Our study demonstrates that SCD5 plays a critical role in GBM where it regulates lipid metabolism, tumor growth, and cell cycle progression. The observed role of SCD5 in cell cycle regulation aligns with previous reports implicating it in cyclin D1–mediated neuronal proliferation ^44^. Notably, SCD5 expression is markedly reduced in differentiated cells and in serum-cultured GBM cell lines, which predominantly rely on SCD1 for fatty acid desaturation. This context-dependent expression underscores the need for careful model selection when investigating SCD5 function. Importantly, our metabolic and genetic analyses reveal functional redundancy between SCD1 and SCD5. In cells with high SCD1 activity, ectopic expression of SCD5 may disrupt lipid homeostasis by driving excessive MUFA production, potentially leading to lipotoxicity or lethal phospholipid imbalances. This mechanistic insight may help reconcile previously conflicting findings regarding the role of SCD5 in various cancer contexts.

Although both SCD1 and SCD5 exhibit delta-9 desaturation activity, their expression patterns, and likely regulatory mechanisms, diverge. SCD1 is broadly expressed and known to be regulated by transcription factors such as SREBP1, LXR, and PPARα, while SCD5 is preferentially expressed in neural tissues and pancreatic cells ^25,45^, suggesting cell type-specific metabolic roles. Notably, a prior study reported that some patient-derived GBM lines with low SCD1 expression can still synthesize MUFA through undetermined mechanisms^18^. Our data now implicate SCD5 as a compensatory enzyme in these contexts. The reciprocal regulation between SCD1 and SCD5 observed in our experiments further highlights their interdependence in maintaining lipid homeostasis.

The pharmacological specificity of existing SCD inhibitors remains unclear, particularly regarding their ability to target both SCD1 and SCD5. Our findings support the development of dual SCD1/SCD5 inhibitors for GBM therapy based on three key observations. First, single-cell RNA sequencing reveals that SCD5 maintains uniform expression across GBM subpopulations, suggesting broad therapeutic relevance. Second, while SCD1 exhibits ubiquitous expression, the brain-enriched distribution of SCD5 offers potential for reduced systemic toxicity, though potential effects on oligodendrocytes and other neural cells require further evaluation. Third, and most critically, SCD5 silencing potently impairs three fundamental GSC properties: self-renewal, tumor initiation, and cell cycle progression, while inducing parthanatos-mediated cell death. This triple vulnerability suggests SCD5 inhibition could simultaneously target multiple oncogenic processes in GBM.

SCD inhibition leads to SFA accumulation causing DNA damage and PARP activation, likely due to elevated levels of reactive oxygen species ^46–48^. Prolonged SFA exposure leads to PARP1 hyperPARylation, AIF nuclear translocation, and ultimately parthanatos. While palmitic acid (C16:0) has been associated with apoptosis ^49^, our data indicate that GSCs, which express high levels of PARP1, can also undergo parthanatos. We propose that the intracellular PARP1 level is a determinant of cell death modality: lower PARP1 expression favors apoptosis, while elevated PARP1 levels shift cell fate toward parthanatos. Importantly, GSCs show enriched PARP1 expression, consistent with previous reports linking PARP1 to stemness maintenance ^50^. This differential vulnerability offers a therapeutic window whereby differentiated tumor cells may undergo apoptosis, while apoptosis-evading GSCs ^51^ could be selectively eliminated through parthanatos.

Standard GBM therapies such as radiation and temozolomide (TMZ) are limited by the robust DNA damage response, particularly trough PARP-dependent repair mechanisms ^52^. Fatty acid oxidation has been shown to fuel PARP activity and support HDR ^53^. Additionally, radiotherapy increases levels of UFAs in GBM, potentially promoting resistance ^54^. Our previous work demonstrated that SCD inhibition reduces RAD51 expression and impairs HDR, sensitizing GBM cells to TMZ and radiation ^20,26,52^, and PARPi-resistant triple-negative breast cancer cells to PARPi ^53^. We now extend these findings by showing that SFA-induced PARP1 depletion downregulates RAD51, providing further support for a mechanistic link between lipid desaturation and DNA repair pathways. These effects parallel findings where PARP1 disruption suppresses RAD51 expression ^35^, and reinforce the bidirectional regulation between lipid metabolism and DNA repair pathways.

Our data support a model in which SFA accumulation induces DNA damage and PARP1 hyperactivation, resulting in depletion of NAD+ and ATP ^55^, and initiating the degradation of PARP1 via lysosomal ^48^ and proteasomal pathways ^56^. This energetic crisis renders GBM cells vulnerable to further stress and DNA damage. Beyond direct tumor cell death, SCD inhibition may also enhance anti-tumor immunity. Damage-associated molecular patterns (DAMPs) release during parthanatos can stimulate immune responses ^57^, as we recently demonstrated in an immunocompetent brain metastasis model where SCD inhibition enhanced dendritic cell activation, interferon signaling, and T-cell-mediated tumor control ^58^. These results suggest that targeting SCD could yield a dual therapeutic benefit. A direct tumor cell killing and immune microenvironment reprogramming, to favor anti-tumor immunity.

### Limitations of the study

While our lipidomic analyses indicate distinct substrate specificities between SCD5 and SCD1 that influence the GBM lipidome, further biochemical characterization of these isoforms is needed to fully understand their functional differences. These findings raise important questions about the physiological and pathological roles of SCD5 in central nervous system disorders, including gliomas, brain metastases, neurodegeneration, and neuroinflammatory conditions. Critical gaps remain in our understanding of SCD5 regulation, particularly regarding its transcriptional and post-translational control mechanisms and how these differ from SCD1. Future studies addressing these questions will be essential to determine potential compensatory interactions between SCD isoforms and whether upregulation of either enzyme could represent a mechanism of acquired resistance to SCD-targeted therapies or DNA-damaging treatments.

In conclusion, our study reveals a mechanistic connection between fatty acid desaturation and DNA repair in GBM, identifying SCD5 as a critical metabolic vulnerability in glioma stem cells. Inhibition of SCD activity leads to saturated fatty acid accumulation, which triggers PARP1 hyperactivation and parthanatos, thereby disrupting tumor maintenance. These effects may be further amplified in combination with DNA-damaging therapies or immunotherapies. Together, our findings support the development of SCD5-specific inhibitors as a promising therapeutic strategy for GBM.

## Materials and methods

### Cell Culture and Differentiation

Primary glioblastoma stem-like cells (GSCs) were obtained from GBM patients at Massachusetts General Hospital (MGG6 and MGG8; provided by H. Wakimoto) under approved IRB protocols, or from I. Nakano (157 and 83) and J. Sarkaria at the Brain Tumor PDX National Resource, Mayo Clinic (M76, M12, and M120). Cells were cultured as neurospheres in DMEM/F12 (Thermofisher, Cat. No. 11330032) supplemented with B27 without vitamin A (1:50; Thermofisher, Cat. No. 12587010), heparin (2 μg/mL; Sigma-Aldrich, 375095-100KU), recombinant human EGF (20 ng/mL; ABM, Cat. No. Z100139), recombinant FGF2 (10 ng/mL; ABM, Cat. No. Z101455), GlutaMAX (1:100; Thermofisher, Cat. No. 35050061), and penicillin/streptomycin (1:100; Thermofisher, Cat. No. 15140122).

Normal human astrocytes (NHAs) were obtained from ScienCell Research Laboratories and cultured in ScienCell astrocyte medium (Catalog No. 1801).

To induce GSC differentiation, cells were dissociated using Accutase (Thermofisher, Cat. No. A1110501) and plated on tissue culture-treated plates in DMEM/F12 supplemented with B27, GlutaMAX, penicillin/streptomycin, and BMP4 (20 ng/mL; BioLegend, Cat. No. 595202) for 6 days. The medium was refreshed after 3 days.

### Chemicals

The following inhibitors and reagents were used in this study: FASN inhibitor (MedChemExpress, GSK2194069) at 50 nM. SCD inhibitor (Cayman Chemical, CAY10566), PARG inhibitor (MedChemExpress, PDD00017273) at 1 and 3 µM, Cycloheximide (Sigma-Aldrich, Cat. No. 01810-1G) at 100 µg/ml, Hydrogen peroxide (Fisher Scientific, Cat. No. H325-100) at 0.5 and 1 mM.

### Lentiviral Production

HEK293T cells (5 × 10⁶) were seeded in 150 mm dishes. After 24 h, 30 µL chloroquine (25 mM; Sigma-Aldrich) was added, and cells were transfected with 15 µg of plasmid encoding the gene or shRNA of interest, 3.75 µg PMD2.G (Addgene, Cat. No. 12259), and 11.25 µg psPAX2 (Addgene Cat. No. 12260) using PEI (Polysciences; 1:3 DNA:PEI ratio).

At 72 h post-transfection, the medium was centrifuged (500 × g, 10 min) and filtered (0.22 µm PES filter: Corning, Cat. No. 431222). The filtrate was ultracentrifuged at 70,000 × g for 90 min at 4°C. Viral pellets were resuspended in 200 µL of 1% BSA in PBS, aliquoted, and stored at −80°C.

### Lentiviral Transduction

GSCs were dissociated with Accutase and seeded at 5 × 10⁵ cells/mL in 12-well plates. Polybrene (1 µg/mL; Millipore, Cat. No. TR-1003-50UL) was added to enhance transduction efficiency. Virus was added and incubated for 8–14 h at 37°C. Media were then replaced, and cells were expanded for subsequent experiments.

### Cell Viability Assay

Cells were plated in 96-well plates in 100 µL total volume. Viability was assessed using the CellTiter-Glo 2.0 Assay (Promega, Cat. No. G9242) according to the manufacturer’s instructions. The reagent was diluted 1:3 in PBS, and 25 µL was added per well. After a 10-minute incubation, 95 µL from each well was transferred to an opaque 96-well plate, and luminescence was measured using a Tecan plate reader.

### Mouse Orthotopic Brain Tumor Models

All animal procedures were approved by the Massachusetts General Hospital Subcommittee on Research Animal Care and followed NIH guidelines. GSCs (10,000 cells) expressing Firefly luciferase (Fluc) were implanted into the left forebrain of athymic nude mice (1.0 mm anterior, 2.0 mm lateral to bregma, 2.5 mm depth) using a stereotactic frame. Tumor growth was monitored by bioluminescence imaging (Xenogen IVIS 200, PerkinElmer) after intraperitoneal injection of D-luciferin (150 mg/kg; Gold Biotechnology, LUCK-100). Signal intensity was quantified using Living Image 4.3.1 software.

### Immunoblotting

Cells were lysed in 1× RIPA buffer (Millipore, Cat. No. 20-188) for 20 minutes on ice, followed by one freeze–thaw cycle. Protein concentration was determined using the BCA assay (Thermo Fisher), and equal amounts of protein were loaded onto NuPAGE 4–12% Bis-Tris gels (Thermofisher). After transfer to PVDF membranes, blots were blocked with 5% skimmed milk in TBST (TBS + 0.5% Tween-20; Sigma-Aldrich, P1379-500ML) and incubated overnight at 4 °C with primary antibodies diluted in 2.5% milk. Membranes were washed and incubated with secondary antibodies for 1 hour at room temperature. Detection was performed using SuperSignal West Pico PLUS or West Femto chemiluminescent substrates (Thermofisher). The complete list of antibodies used for immunoblotting is provided in Supplemental Table 1.

### Immunocytochemistry

Cells were plated on glass coverslips coated with poly-D-lysine (1 h) and laminin (20 µg/mL, overnight; Sigma-Aldrich). After 4 hours, cells were fixed in 4% paraformaldehyde for 20 minutes and blocked with 5% normal goat serum, 0.5% BSA, and 0.1% Triton X-100 in TBS.

Primary antibodies were incubated overnight at 4 °C. After washing, secondary antibodies were applied, and Hoechst (1:1000; Thermofisher) was added during the secondary incubation. Slides were mounted using ProLong Diamond (Thermofisher) and imaged on a Keyence BZ-X810 or Nikon W1-SoRa confocal microscope. The complete list of antibodies used for immunofluorescence is provided in Supplemental Table 2.

### Endoplasmic Reticulum and Mitochondria Imaging

GSCs expressing miRFP670-nano-SCD1 or miRFP670-nano-SCD5 were seeded at 20,000 cells/cm² on laminin-coated (10 µg/mL) glass-bottom 8-well chambers (Ibidi). After 24 h, cells were stained with:ER-Tracker Green (1 µM; Thermofisher) for 30 min at 37°C. MitoView Green (200 nM; Biotium) for 15 min at 37°C. Cells were then imaged using a NIKON W1-SoRa confocal microscope.

### RNA Expression Analysis

Total RNA was extracted using the Quick-RNA Miniprep Kit (Zymo) and quantified with a NanoDrop spectrophotometer (Thermo Fisher). cDNA was synthesized from 600 ng of RNA using the LunaScript RT SuperMix Kit (NEB). Quantitative PCR (qPCR) was performed using the Luna Universal qPCR Master Mix (NEB) on a QuantStudio 3 system (Applied Biosystems). Relative gene expression was calculated using the comparative Ct method (ΔΔCt), with normalization to TBP as a housekeeping gene. The list of primers used in this study is provided in Supplemental Table 3.

### Bulk RNA Sequencing

Total RNA (500 ng) was extracted with the Quick-RNA Miniprep Kit and assessed for quality on a Bioanalyzer (Agilent). Libraries were prepared using the NEBNext Ultra II Directional RNA Library Prep Kit (NEB), with poly(A) selection, fragmentation, cDNA synthesis, end repair, adapter ligation, and PCR amplification. Libraries were sequenced on an Illumina NextSeq 2000 (paired-end).

Reads were aligned to GRCh38 using the Rsubread package in R. Gene counts were generated using featureCounts, and differential expression was analyzed with DESeq2. Significance was defined as adjusted p < 0.05.

### Shotgun Lipidomics

Cells were transferred into extraction tubes containing phosphate-buffered saline (PBS). Lipid extraction was performed using a modified Bligh and Dyer protocol as previously described ^59^. Prior to the biphasic extraction step, each sample was spiked with a mixture of 74 lipid standards (Avanti: 330820, 861809, 330729, 330727, 791642, 330726). Following two sequential extractions, the combined organic phases were dried using a Thermo SpeedVac SPD300DDA set to ramp setting 4 at 35 °C for 45 minutes, with a total run time of 90 minutes.

Dried lipid extracts were resuspended in a 1:1 mixture of methanol and dichloromethane containing 10 mM ammonium acetate and transferred to robovials (Thermofisher, Cat. No. 10800107) for analysis. Samples were analyzed by direct infusion using a Sciex 5500 mass spectrometer equipped with a Differential Mobility Device (DMS). Targeted acquisition included 1,450 lipid species spanning 17 subclasses. The DMS was tuned using the EquiSPLASH LIPIDOMIX standard (Avanti: 330731). Data processing was performed using an in-house workflow. Instrument parameters, MRM transitions, and the analytical method are described in Su B. (2021)^60^. Quantification was normalized to cell number.

### Cell Cycle Synchronization

GSCs were synchronized at the G1/S phase using 2 mM thymidine (Sigma-Aldrich, T1895-1G). Cells were treated for 16 h, released into thymidine-free medium for 8 h, then treated again for 16 h. After the second treatment, cells were washed and placed in fresh medium. This time point was designated as time 0.

### Flow Cytometry for Cell Cycle Analysis

GSCs were transduced with SCD1/5-targeting shRNA and cultured for 4 days. Cells were fixed in 70% ethanol after PBS washes. For cell cycle profiling, cells were stained with propidium iodide (0.02 mg/mL), RNase A (0.5 mg/mL; NEB), and Triton X-100 (0.1%) in PBS, incubated at 37°C for 30 min, and analyzed using a Cytek Aurora flow cytometer.

### Statistical Analysis

All statistical analyses were performed using R v4.3.0. Each experiment included at least four biological replicates. Statistical comparisons were made using two-tailed Student’s t-tests. Data normality was assessed using the Shapiro-Wilk test. Non-parametric tests were used when data were non-normal or heteroscedastic. P < 0.05 was considered statistically significant.

## Acknowledgments

This work was supported by NIH/NINDS R01 NS113822 (C.E.B.), DoD Peer Reviewed Cancer Research CA191075 (C.E.B.), NIH/NCI P50 CA165962 SPORE in Brain Tumor Research subaward (C.E.B.).The authors thank the MGH NextGen Sequencing Core (Boston, MA) for sequencing services and the Mass General Brigham Center of Excellence for Molecular Imaging (Boston, MA) for their support and resources.

