## Supplemental information for "SCD1 and SCD5 Modulate PARP-Dependent DNA Repair via Fatty Acid Desaturation in Glioblastoma"

**Supplemental Table 1:** List of antibodies used for Immunoblot.

| Target | Brand | Reference |
|---|---|---|
| SCD5 | Thermofisher | PA5-89006 |
| SCD5 | SynoBiologics | Custom maid |
| SCD1 | Thermofisher | MA5-27542 |
| PARP1 | Thermofisher | 14-6667-82 |
| PAR | Cell Signaling Technologies | 89190S |
| γH2A.X | Cell Signaling Technologies | 9718S |
| AIF | Cell Signaling Technologies | 5318S |
| LC3 A/B | Cell Signaling Technologies | 12741S |
| GFP | Thermofisher | MA5-15256 |
| TBP | Proteintech | 66166-1-Ig |
| β-Tubulin | Cell Signaling Technologies | 86298S |
| β-Actin | Cell Signaling Technologies | 3700S |
| GAPDH | Proteintech | 60004-1-Ig |
| Anti-rabbit HRP | Cell Signaling Technologies | 7074V |
| Anti-mouse HRP | Cell Signaling Technologies | 7076V |

**Supplemental Table 2:** List of antibodies used for immunofluorescence or immunohistochemistry.

| Target | Brand | Reference |
|---|---|---|
| SCD5 | SynoBiologics | Custom maid |
| SCD1 | Thermofisher | MA5-27542 |
| PAR | Cell Signaling Technologies | 89190S |
| γH2A.X | Cell Signaling Technologies | 9718S |
| Goat anti-rabbit AF 647 | Thermofisher | A-21245 |
| Goat anti-mouse AF 488 | Thermofisher | A-11001 |
| Goat anti-rabbit Biotin-XX | Thermofisher | B-2770 |
| Goat anti-mouse Biotin-XX | Thermofisher | B-2763 |

**Supplemental Table 3:** Primers used for RT-PCR.

| Target | Forward primer | Reverse primer |
|---|---|---|
| *SCD5* | CACTCTGCTCTGGGCCTACTT | AGTTGGCGACAGCCAGAAATA |
| *SCD1* | TCTAGCTCCTATACCACCACCA | TCGTCTCCAACTTATCTCCTCC |
| *PARP* | GAGGTGGATGGGTTCTCTGA | ACACCCCTTGCACGTACTTC |
| *SOX2* | AACCCCAAGATGCACAACTC | GCTTAGCCTCGTCGATGAAC |
| *NESTIN* | AACAGCGACGGAGGTCTCTA | TTCTCTTGTCCCGCAGACTT |
| *GFAP* | AGGAAGATTGAGTCGCTGGA | ATACTGCGTGCGGATCTCTT |
| *OLIG1* | CCCCAAAAGTAGCGTAACCA | CCGGTACTCCTGCGTGTTA |
| *OLIG2* | CTCCTCAAATCGCATCCAGA | CTCCTCAAATCGCATCCAGA |
| *TBP* | CCACTCACAGACTCTCACAAC | CTGCGGTACAATCCCAGAACT |

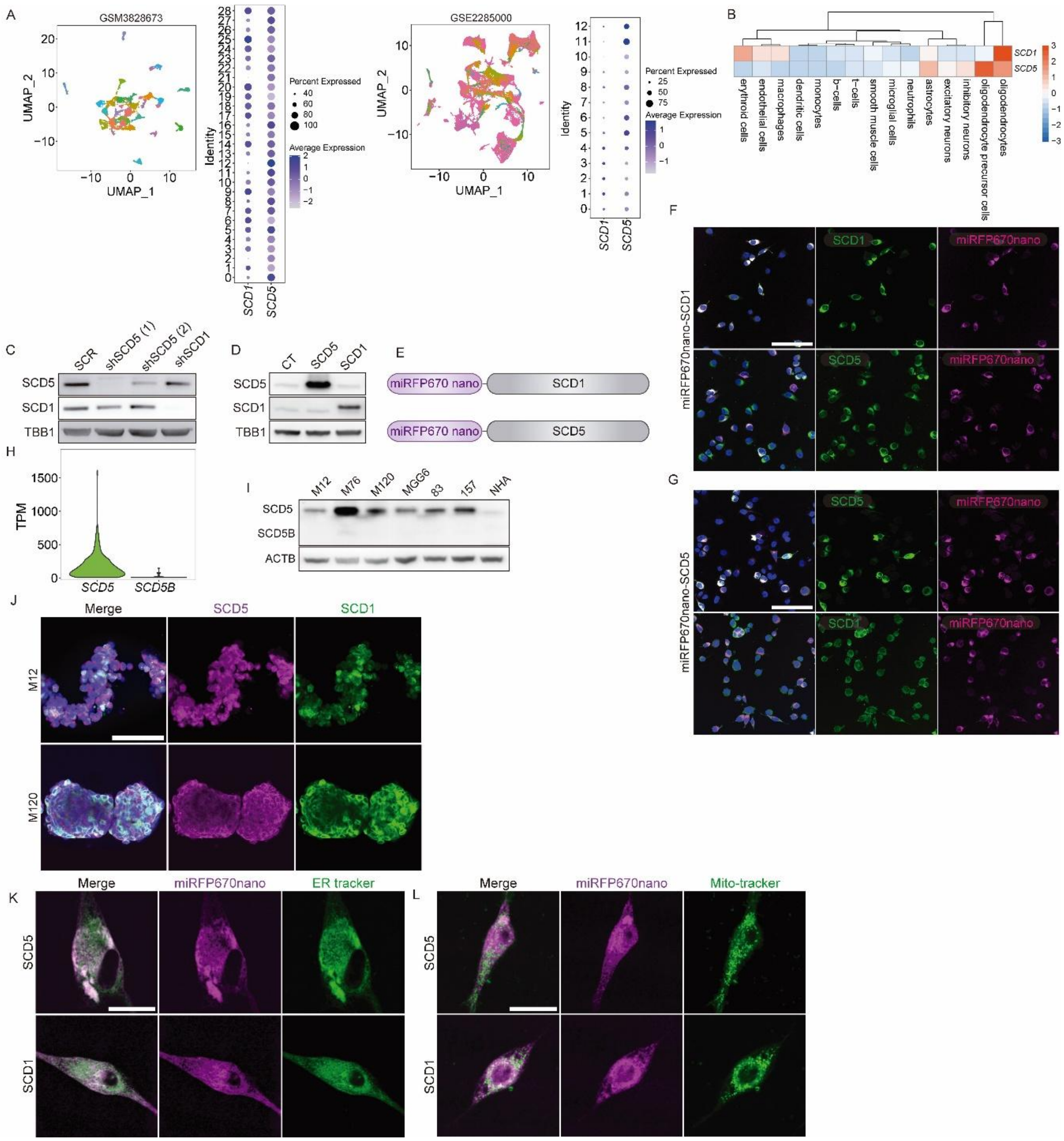

**Supplemental Figure 1. SCD5 expression profile in GBM cells**

(A) UMAP projection and dot plot of SCD1/SCD5 expression across cellular subpopulations in two GBM patient scRNA-seq datasets (GSM3828673, GSE2285000).

(B) Heatmap of SCD1/SCD5 expression in normal brain cell types (HPA scRNA-seq data).

(C-D) Immunoblot analysis of SCD expression after SCD1/SCD5 knockdown (for 4 days; C) or overexpression (D).

(E) Schematic of C-terminally tagged SCD1/SCD5-miRFP670nano fusion proteins.

(F-G) Immunofluorescence localization of (F) SCD1-miRFP670nano and (G) SCD5-miRFP670nano in U251 cells. Scale bars: 40 μm.

(H) Bulk RNA-seq analysis of SCD5/SCD5B isoform expression in glioma (GLASS consortium data).

(I) Immunoblot detection of SCD5 (~35 kDa) and SCD5B (~29 kDa) isoforms in GSCs and NHA.

(J) Immunostaining of endogenous SCD1/SCD5 in GSCs detected by confocal microscopy. Scale bar: 50 μm.

(K-L) Live-cell confocal imaging of (K) ER (ER-Tracker Green) and (L) mitochondrial (MitoTracker Green) co-localization with SCD1/SCD5-miRFP670nano in U251 cells. Scale bars: 25 μm.

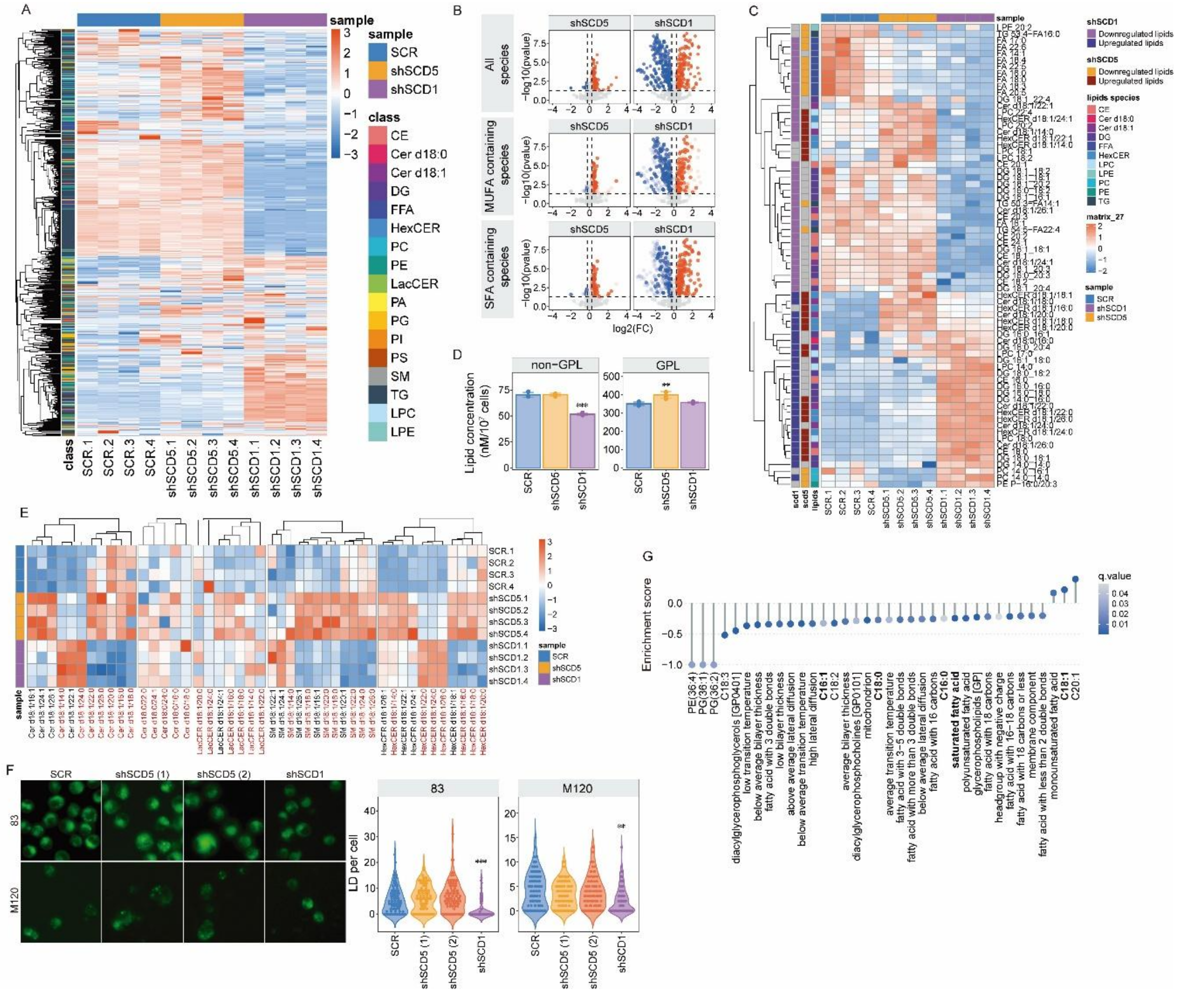


**Supplemental Figure 2. SCD1 and SCD5 knockdown induce distinct lipidomic alterations in GSCs.**
(A) Heatmap of global lipid profiles in GSCs following 4-day SCD1 or SCD5 knockdown.
(B) Volcano plots of lipid species changes (top: all lipids; middle: MUFA-containing species; bottom: SFA-containing species). Filled dots indicate significant changes (p<0.05).
(C) Heatmap of the top 25 most significantly altered lipid species post-knockdown.
(D) Quantification of glycerophospholipid (GPL) and non-GPL levels in knockdown conditions.
(E) Sphingolipid subspecies heatmap, with saturated acyl chain-containing lipids highlighted in red.
(F) BODIPY staining (left) and quantification (right) of lipid droplet accumulation (n>100 cells; *p<0.05, **p<0.01, ***p<0.001, Student's t-test compared to SCR).
(G) Triglyceride carbon number distribution analysis in SCD1- versus SCD5-deficient GSCs.
(H) LION enrichment analysis of lipid-associated pathways in SCD5-overexpressing GSCs (n=4).

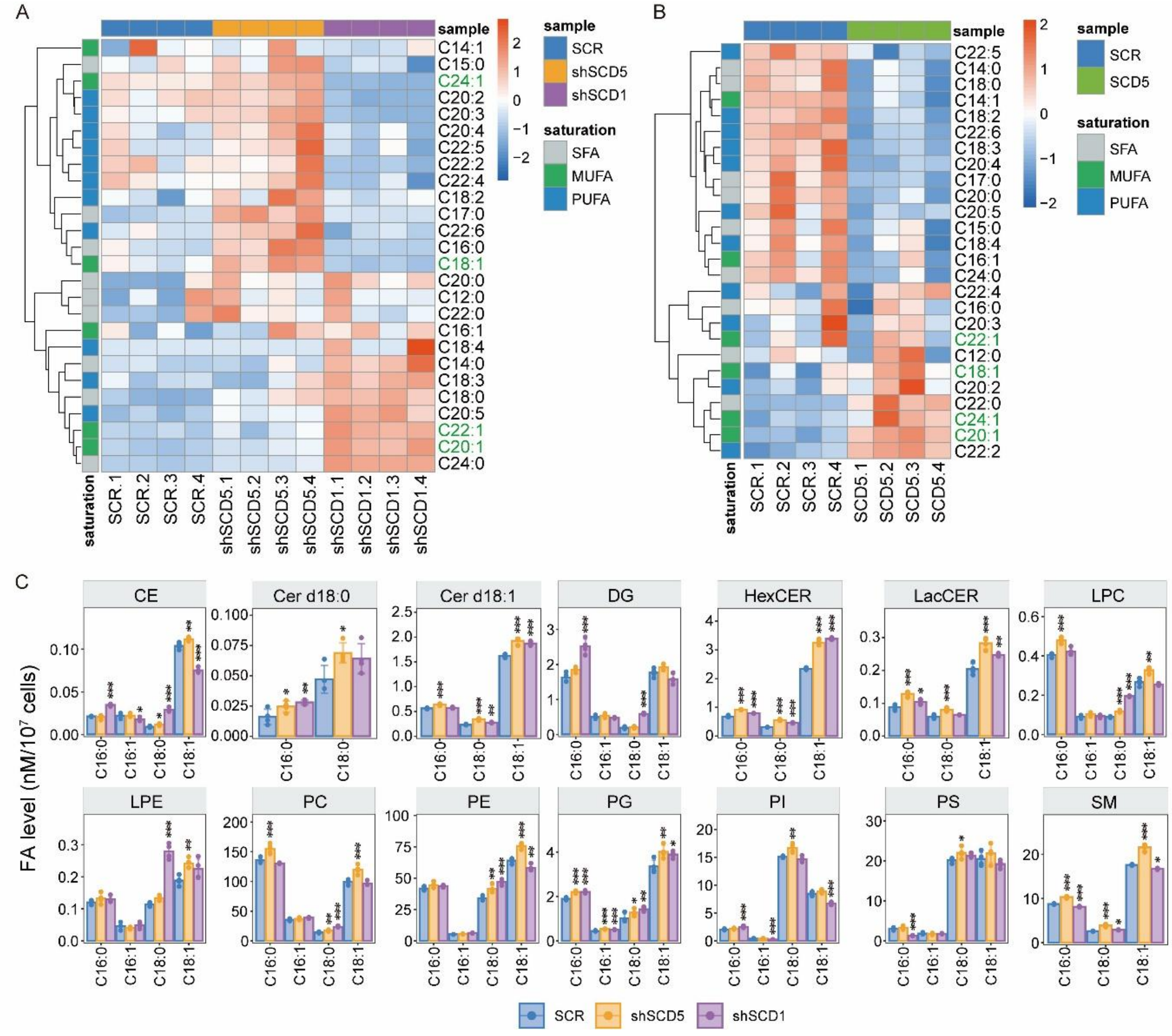


**Supplemental Figure 3. SCD5 exhibits substrate preference for 18:0 over 16:0 in GSCs.**

(A) Fatty acid composition heatmap from shotgun lipidomics of SCD1- or SCD5-knockdown (KD) GSCs. Fatty acids are categorized as saturated (SFA), monounsaturated (MUFA), or polyunsaturated (PUFA), with ≥18-carbon MUFAs highlighted in green.

(B) Corresponding fatty acid composition heatmap in SCD5-overexpressing GSCs (classification as in A).

(C) Quantification of lipid class-specific SCDs substrates (C16:0 and C18:0 ) and products (C16:1 and C18:1) following 4-day SCD5 knockdown (mean ± SD; n=4; *p<0.05, **p<0.01, ***p<0.001, Student's t-test).

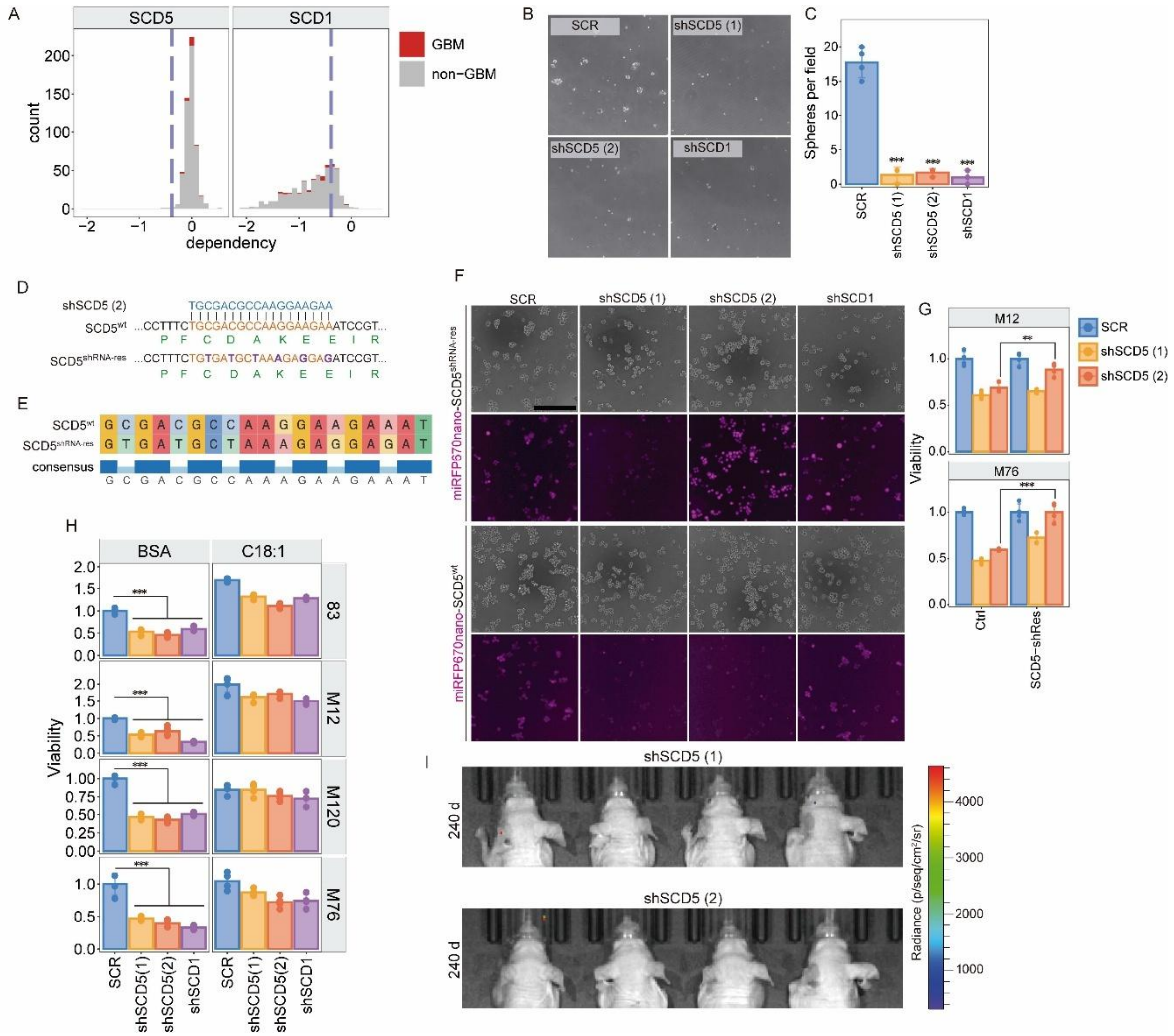

**Supplemental Figure 4. SCD1 and SCD5 are required for GSC maintenance and tumorigenesis.**

(A) Dependency scores for SCD1 and SCD5 across cancer cell lines from the DepMap database. GBM cell lines are highlighted in red

(B) Secondary neurosphere formation in SCD1- or SCD5-knockdown GSCs (M120 line, 4-day after transduction).

(C) Quantification of neurospheres per field in the secondary neurosphere formation assay (mean ± SD).

(D) Engineering strategy for shRNA-resistant SCD5 open reading frame (SCD5$^{shRNA\text{-}res}$), incorporating six silent mutations.

(E) Next-generation sequencing validation of the SCD5$^{shRNA\text{-}res}$ confirming the introduced mutations in the shRNA-resistant SCD5 plasmid.

(F) Fluorescence imaging of GSCs expressing miRFP670nano-tagged SCD5 or SCD5$^{shRNA\text{-}res}$ post-shSCD5 knockdown. Scale bar: 100 µm.

(G) Cell viability rescue in SCD5$^{shRNA\text{-}res}$-expressing GSCs after SCD5 knockdown (mean ± SD; n=4; *p<0.05, **p<0.01, ***p<0.001, Student's t-test).

(H) Cell viability of SCD1/SCD5-knockdown GSCs supplemented with C18:1 at 50 µM (mean ± SD; n=4).

(I) Tumor-free survival in mice implanted with SCD1/SCD5-deficient GSCs (240 days post-implantation; controls succumbed by day 60).

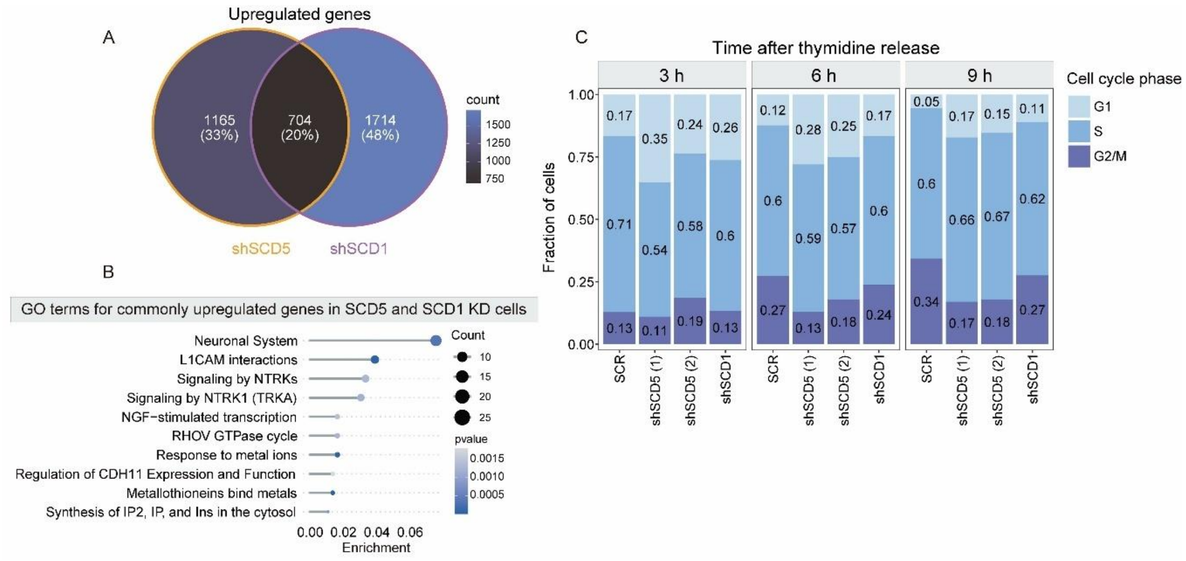


**Supplemental Figure 5. SCD1 and SCD5 regulate cell cycle progression in GSCs.**
(A) Venn diagram of downregulated transcripts shared between SCD1- and SCD5-deficient GSCs.
(B) GO enrichment analysis of commonly upregulated genes in SCD1/SCD5 knockdown conditions.
(C) Cell cycle distribution analysis by flow cytometry of GSCs synchronized with thymidine and released for 3, 6, or 9 hours post-SCD1/SCD5 knockdown.

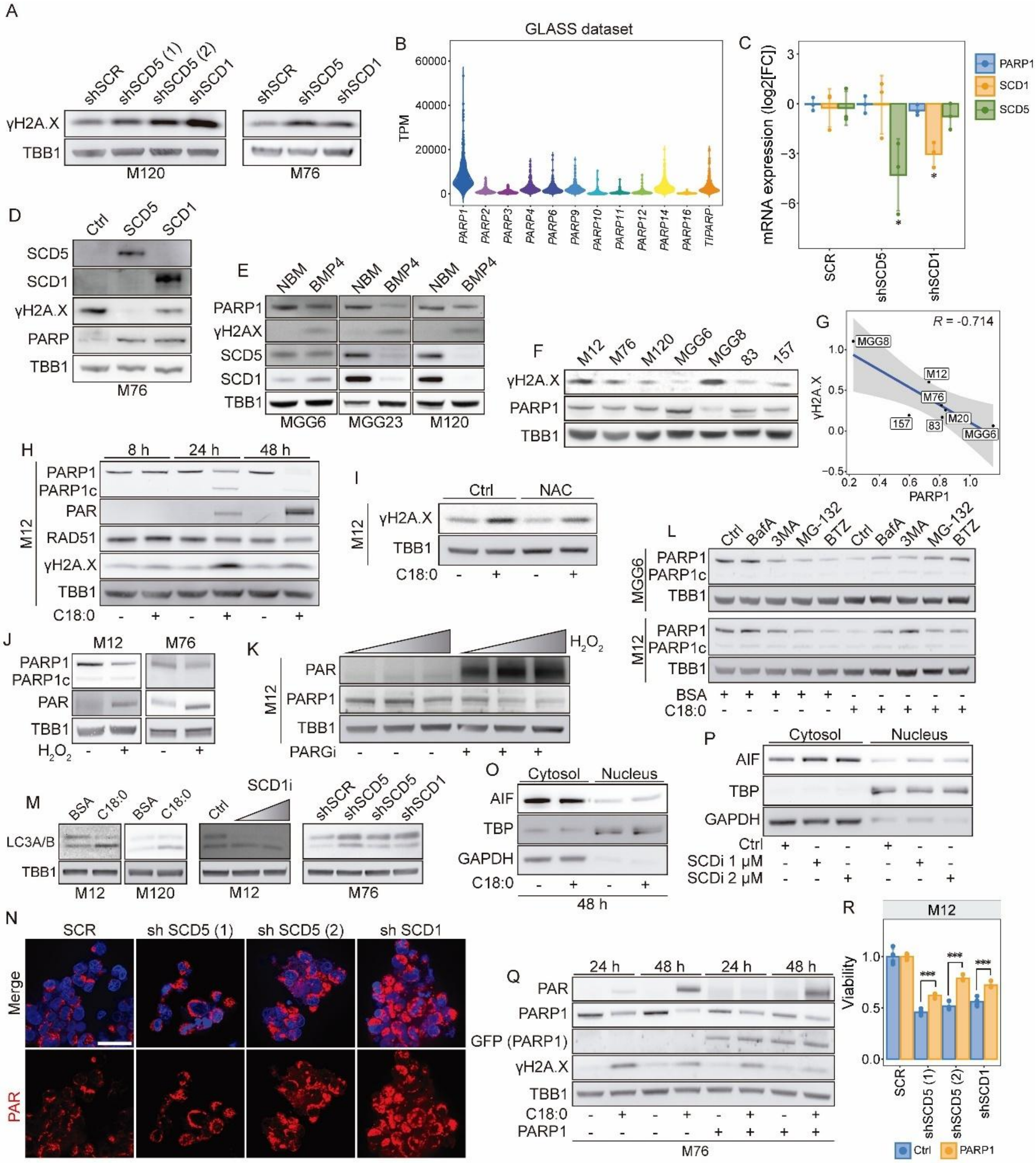
A
shSCR
shSCD5 (1)
shSCD5 (2)
shSCD1
γH2A.X
TBB1
M120
shSCR
shSCD5
shSCD1
M76
B
GLASS dataset
TPM
60000
40000
20000
0
PARP1
PARP2
PARP3
PARP4
PARP6
PARP9
PARP10
PARP11
PARP12
PARP14
PARP16
TIPARP
C
mRNA expression (log2[FC])
3
0
-3
-6
SCR
shSCD5
shSCD1
PARP1
SCD1
SCD5
D
Ctrl
SCD5
SCD1
SCD5
SCD1
γH2A.X
PARP
TBB1
M76
E
NBM
BMP4
PARP1
γH2AX
SCD5
SCD1
TBB1
MGG6
MGG23
M120
F
M12
M76
M120
MGG6
MGG8
83
157
γH2A.X
PARP1
TBB1
G
R = -0.714
γH2A.X
PARP1
MGG8
M12
M76
M20
157
83
MGG6
H
8 h
24 h
48 h
M12
PARP1
PARP1c
PAR
RAD51
γH2A.X
TBB1
C18:0
I
Ctrl
NAC
M12
γH2A.X
TBB1
C18:0
J
M12
M76
PARP1
PARP1c
PAR
TBB1
H2O2
K
H2O2
M12
PAR
PARP1
TBB1
PARGi
L
Ctrl
BafA
3MA
MG-132
BTZ
MGG6
PARP1
PARP1c
TBB1
M12
BSA
C18:0
M
BSA
C18:0
SCD1i
Ctrl
shSCR
shSCD5
shSCD5
shSCD1
LC3A/B
TBB1
M12
M120
M76
O
Cytosol
Nucleus
AIF
TBP
GAPDH
C18:0
48 h
P
Cytosol
Nucleus
AIF
TBP
GAPDH
Ctrl
SCDi 1 μM
SCDi 2 μM
N
SCR
sh SCD5 (1)
sh SCD5 (2)
sh SCD1
Merge
PAR
Q
24 h
48 h
PAR
PARP1
GFP (PARP1)
γH2A.X
TBB1
C18:0
PARP1
M76
R
M12
Viability
1.0
0.5
0.0
***
SCR
shSCD5 (1)
shSCD5 (2)
shSCD1
Ctrl
PARP1

**Supplemental Figure 6. SFA accumulation induces DNA damage, PARP1 hyperactivation, and parthanatos in GSCs.**

(A) Immunoblot analysis of SCD1, SCD5, and γH2AX in GSCs following 4-day SCD1/SCD5 knockdown.

(B) PARP family member expression in GBM tumors from the GLASS consortium bulk RNA-seq database.

(C) qPCR analysis of PARP1 mRNA expression in three GSC lines (M12, M76, and M120) 4 days post-SCD1/SCD5 knockdown (mean ± SD; n=3).

(D) Immunoblot analysis of SCD1, SCD5, PARP1, and γH2AX in SCD1/SCD5-overexpressing GSCs.

(E-F) PARP1-γH2AX correlation analysis in patient-derived GSCs (F: Spearman's R).

(G) Immunoblot analysis for PARP1 and γH2AX levels in BMP4-differentiated GSCs (6 days). Please note that this is the same blot used to detect SCD1 and SCD5 levels in Figure 2E.

(H) Immunoblot analysis of the indicated proteins following treatment with C18:0 (150 µM for 8, 24 and 48 hours).

(I) Immunoblot analysis of γH2A.X in GSCs pre-treated with N-acetylcysteine (NAC, 0.25 mM) for 4 h and then co-treated with C18:0 for 24 h.

(J) Immunoblot analysis of PARP1 and PAR (at the expected molecular weight of PARP1) in GSCs treated with $H_2O_2$ (1 mM) for 15 minutes.

(K) PARP1 hyperPARylation induced by $H_2O_2$ (0-2 mM, 15 min) ± PARG inhibitor (1 µM, 24 h).

(L) Immunoblot analysis of PARP1 in GSCs treated with C18:0 (150 µM) for 24 h, followed by treatment with autophagy inhibitors Bafilomycin A (BafA, 1 µM) and 3-Methyladenine (3MA, 3 mM), and proteasome inhibitors MG132 (2.5 µM) and Bortezomib (BTZ, 250 nM) for 6 h.

(M) Cleaved LC3 levels in GSCs treated with C18:0 (150 µM) for 24 h, SCDi (CAY10566; 0.5 and 1 µM), or after knockdown of SCD1 and SCD5 for 4 days.

(N) PAR immunofluorescence in SCD1/SCD5-knockdown in M12 GSCs for 4 days. Scale bar: 25 µm.

(O-P) Nuclear AIF translocation in GSCs treated with C18:0 (O; 150 µM for 48h) or SCDi (P; 72h).

(Q) Immunoblot analysis of PARP1, PAR, γH2A.X, and GFP in PARP1-GFP-expressing GSCs (M76) treated with C18:0 (150 µM) for 24 and 48 hours.

(R) Cell viability of SCD1/SCD5-knockdown GSCs ± PARP1 overexpression (mean ± SD; n=4; *p<0.05, **p<0.01, ***p<0.001, Student's t-test).
